# Transposable elements impact the human regulatory landscape through cell type specific epigenomic associations

**DOI:** 10.1101/2024.08.07.606967

**Authors:** Jeffrey Hyacinthe, Guillaume Bourque

**Affiliations:** Quantitative Life Sciences, McGill University, Montréal, QC, Canada; Department of Human Genetics, McGill University, Montréal, QC, Canada; Canadian Center for Computational Genomics, McGill University, Montréal, QC, Canada; Victor Phillip Dahdaleh Institute of Genomic Medicine at McGill University, Montréal, QC, Canada

## Abstract

Transposable elements (TEs) are DNA sequences able to create copies of themselves within the genome. Despite their limited expression due to silencing, TEs still manage to impact the host genome. For instance, some TEs have been shown to act as cis-regulatory elements and be co-opted in the human genome. This highlights that the contributions of TEs to the host might come from their relationship with the epigenome rather than their expression. However, a systematic analysis that relates TEs in the human genome directly with chromatin histone marks across distinct cell types remains lacking. Here we leverage a new dataset from the International Human Epigenome Consortium with 4867 uniformly processed ChIP-seq experiments for 6 histone marks across 175 annotated cell labels and show that TEs have drastically different enrichments levels across marks. Overall, we find that TEs are generally depleted in H3K9me3 histone modification, except for L1s, while MIRs were highly enriched in H3K4me1, H3K27ac and H3K27me3 and Alus were enriched in H3K36me3. Furthermore, we present a generalised profile of the relationship between TEs enrichment and TE age which reveals a few TE families (Alu, MIR, L2) as diverging from expected dynamics. We also find significant differences in TE enrichment between cell types and that in 20% of the cases, these enrichments were cell-type specific. Moreover, we report that at least 4% of cell types-histone-TE combinations featured significant differences in enrichment between healthy and cancer samples. Notably, we identify 456 cell type-histone-TE triplets with strong cell-type specific enrichments. We show that many of these triplets are associated with relevant biological processes and genes expressed in the relevant cell type. These results further support a role for TE in genome regulation and highlight novel associations between TEs and histone marks across cell types.

## Introduction

Transposable elements (TEs) are DNA sequences with the ability to duplicate themselves within the genome. This transposition is done through 2 main mechanisms which define TE classes. Retrotransposons are sequences that use an RNA intermediate which reverse transcribes back into DNA and integrates in another genomic location akin to a copy and paste approach; while DNA transposons excise themselves before inserting elsewhere using a cut and paste mechanism^1,2^. Within classes, TEs are grouped into families, which capture the elements origin and history, and also into subfamilies representing more closely related elements and finer divergence within those families^3–5^. Through their replication, TEs have proliferated through genomes and currently cover at least 50% of the human genome^6^.

One reason why TEs have gathered increased attention is due to their involvement in host gene regulation. Indeed, while most TEs in the human genome have lost their transposition ability, some TEs have been found to be associated with enhancers^7^. For instance, they were shown to be associated with the core regulatory network of human embryonic stem cells^8,9^ and some TEs have been co-opted into the human genome such as an ERV by the Aim2 gene^10^. Furthermore, about 750 SVAs were found to act as enhancers or promoters modulating gene expressions in pluripotent cells^11^.

A recent investigation with 19 different cell types showed that TE encompassed one quarter of the regulatory epigenome and that 47% of TE instances could be found in an active state^12^. This study, primarily centered on epigenetic states, highlighted that cell type specific TE associations could be detected. In other studies, the histone mark H3K9me3, a hallmark of heterochromatin, has been shown to be associated with repeat elements^13,14^. More targeted investigations of specific TEs such as Alus or L1 and histone marks have demonstrated that TEs can behave as enhancers^7,15^ and in a cell-type specific manner^16^. These results demonstrate that cell type differentiation and function may be partly regulated by TEs. However, a comprehensive analysis that relates multiple TEs with reference histone marks across distinct cell types is lacking. Thus, we set out to investigate TE’s associations with histone marks to better understand TEs and their relationship with the epigenome.

With the generation of a large epigenome dataset from the International Human Epigenome Consortium (IHEC)^17^, it is now possible to investigate all TEs in the human genome, across reference histone marks in various cell types. The latest IHEC reprocessing EpiATLAS dataset makes available 4867 Chromatin Immunoprecipitation sequencing (ChIP-Seq) datasets from 6 histone marks (H3K4me1, H3K36me3, H3K9me3, H3K27me3, H3K4me3 and H3K27ac), from 175 annotated cell labels which were grouped into 47 cell types^18^. This dataset contains novel cells types for TE investigation, such as lymphocytes of B lineage and thyroid, has more samples of tissues previously reported to be associated with TEs such as twice the brain sample count from NIH Roadmap^19^, has a larger set of repressive mark data to contrast activating marks, contains healthy and disease samples from the same tissue and includes many replicates to characterise the variability of our observations.

Here, we present an overview of TE overlap found in the EpiATLAS dataset and a comprehensive map of TE subfamily enrichment across cell type and histone marks. We investigated the relationship between TE enrichment and TE age, we surveyed the TE enrichment across cell types and whether they changed depending on health status. We measured the cell type specificity of the TE enrichments and identified TE-cell type regulatory candidates in terms of specific and extreme associations. Some candidates were associated with relevant biological processes and reports of genes expression in the given cell type. Taken together, these results provide a consensus resource of the TE profile across cell types and histone marks.

## Results

### A large and varied comprehensive inter-consortium dataset

Our analysis leveraged the EpiATLAS dataset, a large uniformly processed dataset generated by a consortium of consortiums^17,18^. Specifically, we obtained 4867 ChIP-seq samples for 6 histone marks (H3K4me1, H3K36me3, H3K9me3, H3K27me3, H3K4me3 and H3K27ac), coming from 7 consortiums (Blueprint, CEEHRC, DEEP, AMED-CREST, NIH Roadmap Epigenomics, GIS, EpiHK) and prepared by the IHEC EpiATLAS integrative analysis group (Methods). Of the consortiums from which the data originated Blueprint & CEEHRC accounted for more than 2/3 of the samples (Fig 1A). Our dataset consisted of 175 different cell labels which were grouped into 47 broad cell types (Fig 1B). Compared to the NIH Roadmap reference epigenomes^19^, this represents more than five times the number of samples (4867 vs 733), including 6 times the brain samples (476 vs 72) and introduced sample in novel cell types such as various lymphocytes (373) (Fig 1B). It is also more than 3 times the number of samples from the more recent ENCODE epigenome dataset^20^ (See Methods). In IHEC, each cell types contained on average 104 samples with a mean of 18 biological or technical replicates per assay. While most samples did not have all 6 histone marks, they were represented in comparable amounts (Fig 1C) with a larger number coming from H3K27ac (1481). Notably, the EpiATLAS dataset also included samples with different health conditions: 3557 healthy samples, 1007 cancer samples and 303 diseased samples (Fig 1D).

**Figure 1.**
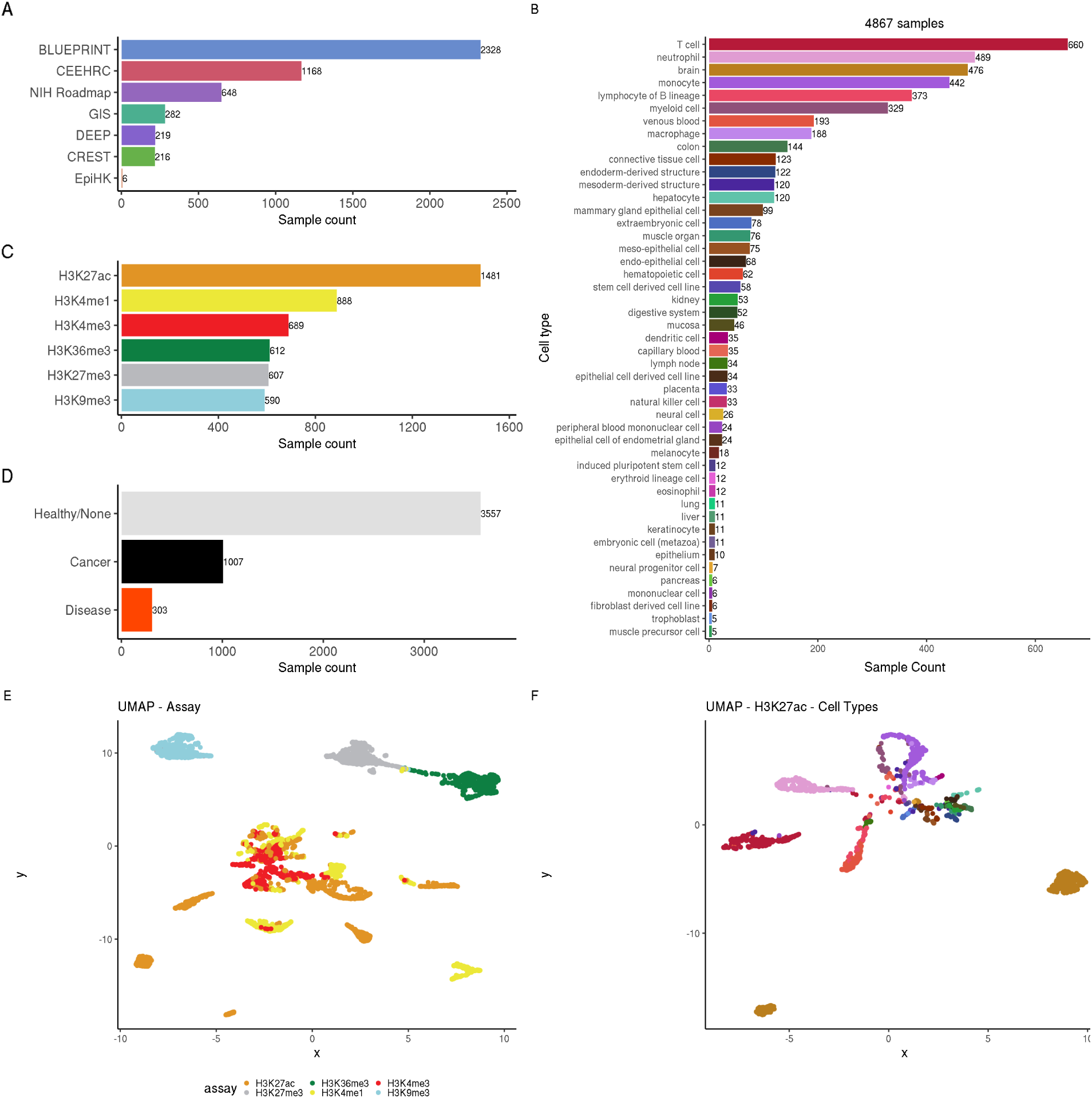
An expansive dataset obtained from IHEC. **A|** Sample count by consortium **B|** Sample count for each cell type **C|** Proportion of samples in from the various assays. **D|**By health status **E|** UMAP on peak counts within 10kb regions across the genome for the 4614 samples. Only the 20,000 windows with most variance (excluding the first 1000) were used. Color represents the assay. **F|** Same as E but only for H3K27ac with cell type coloring from B.

To make sure that the samples obtained from different consortiums were of good quality and comparable, we looked at their distribution of peaks relative to genes transcription start sites (TSS) and found that it remained consistent within assay across consortium (Supplemental Fig 1A). For instance, we find that 84.7% of H3K36me3 peaks were in intragenic regions across all consortiums. This was much lower for H3K9me3 samples at around 33.4%. The distinct distributions between assays combined with consistency across consortium indicated the data captured the similarly located regions across samples. Of the 4867 samples only 253 samples were deemed outliers based on peak distribution and were discarded (Supplemental Fig 1B, Methods). We visualised the 4614 retained samples through a UMAP dimension reduction on the peak counts within genome windows of 10 kilo base pairs (kbp) and found that, as expected, the assay type showed strongest clusters (Fig 1E, Methods). Within the assays, the consortium mostly mixed, but there were some consortium-specific clusters suggesting either cell-type or sample preparation effects (Supplemental Fig 2A). We also note that at that level, cell type associations were not clear, except within some H3K27ac and H3K4me1 datasets (Fig 1E-F, Supplemental Fig 2E). We used PCA as an alternative data visualization method and also found that assays form clusters based on first 2 PCs, while cell types and consortium do not (Supplemental Fig 2B-C,F). The assay clusters are still visible between PC 2 and 3 (Supplemental Fig 2D).

Thus we make use of a more comprehensive dataset, which includes many replicates, novel healthy and disease tissues and is consistent even across consortiums, enabling our TE analysis to explore new associations and patterns.

### TEs have distinct epigenetic marks association profiles

Next, to characterize the contribution of TEs to ChIP-seq peaks, we measured the percentage of peaks within each sample that overlapped such elements. We found that the overlap with TE families remained largely consistent within assays (Fig 2A). On average, 55.8% of peaks were found to overlap with TEs and we observed that the activating mark H3K4me1 (54.2%) and H3K27ac (50.7%) displayed more similar TE overlap profiles. In contrast, the repressing mark H3K9me3 had a distinctively higher TE overlap (78.3%), which is consistent with the reported role of H3K9me3 in TE repression^14,21,22^. We also found that most of the TE overlap came from the top 5 most common families (Alu, L1, MIR, L2, ERVL-MaLR, Supplemental Fig 3A). TEs can cause multi-mapped reads which can lead to analysis issues. To make sure that mappability was not impacting these observations, we used the mappability track from the UCSC Genome browser^23^ (Methods). We found that mappability did not correlate with the overlap we observed and only Telo, SVA, Satellite, Centr and acro families featured a median mappability below 80% (Supplemental Fig 3A-B, 4). Next, we explored the cumulative and mean TE Length and the TE instance count and found that the TE overlap profile was most similar to the instance count (Supplemental Fig 3A, C-E).

**Figure 2.**
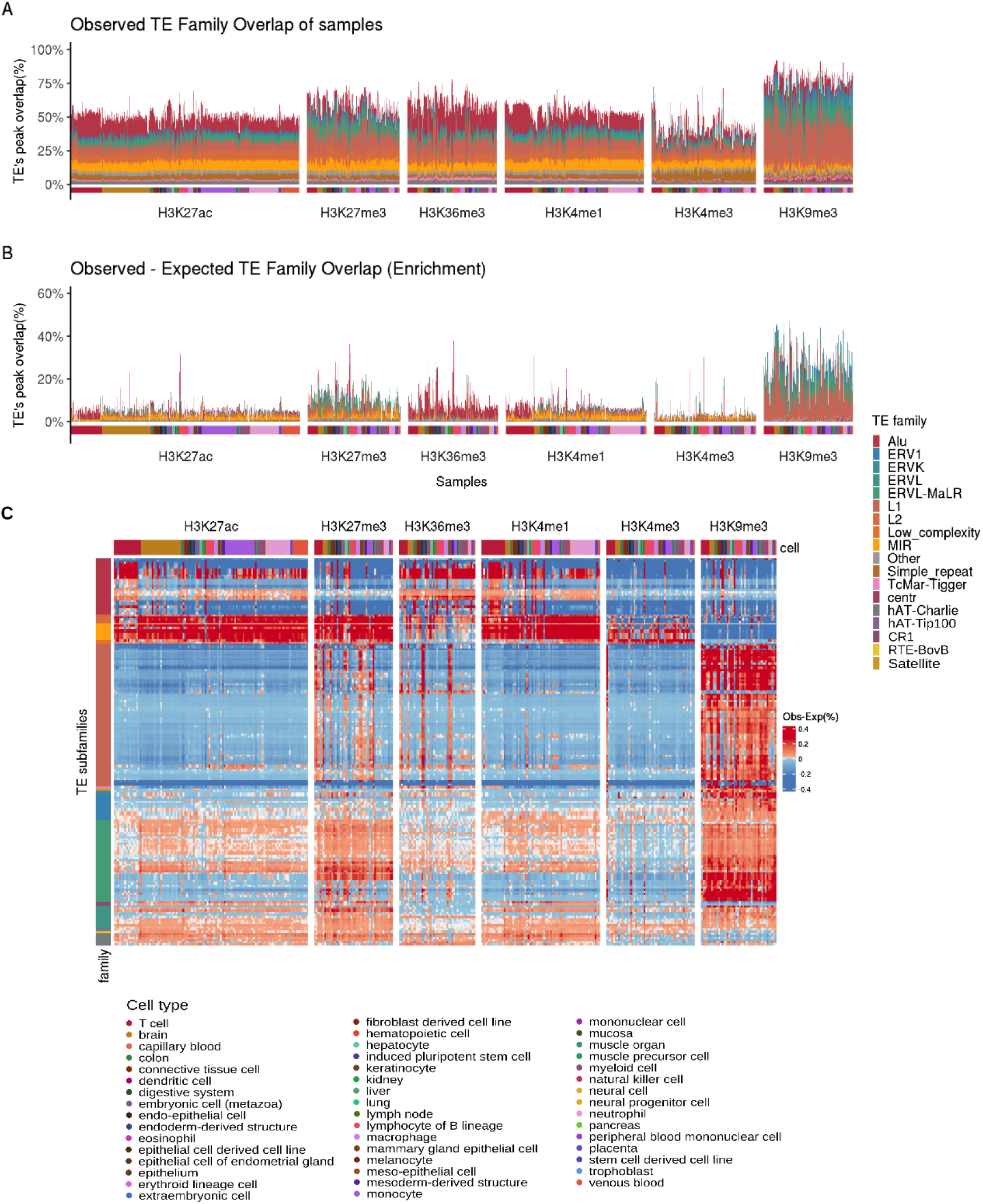
TE displays distinct association profiles with histone modifications. **A|** Percentage of peaks overlapping TEs colored by TE family for the 4614 samples. Samples are annotated by cell type. **B|** Difference between the Observed overlap (A) and the simulated background overlap for using only significantly enriched (p-val < 0.005) TEs. Same annotations as A. **C|** Heatmap of the TE subfamily enrichment of the selected 164 subfamilies for the 4614 samples. Enrichment (obs-exp in percent, including non-significantly enriched TEs). TE subfamilies are grouped by family (x axis), Samples are grouped by Assay (y axis) and annotated for cell type. The samples within groups are hierarchically clustered.

To better capture TE enrichment, we identified TE families that featured significantly higher overlap (p-val < 0.005) than in distribution matched random simulations (Methods). Using the difference between the observed and the expected (obs-exp, which we called an enrichment), we noted distinct patterns such as an enrichment of L1 and ERV TEs in H3K9me3 and Alu in H3K36me3 (Fig 2B). The low enrichment also exposed that while there was high overlap with TE families in the genome, most of it could be attributed to chance and that significant TE associations arose from a minority of regions.

Next, to detect more specific associations, we repeated this enrichment analysis using TE subfamilies instead of families. We found that the majority of TE subfamilies were not significantly enriched (Supplemental Fig 5A) and decided to focused on the most globally enriched subfamilies. This was done by adding up all significant subfamily enrichment across all samples and selecting the 164 subfamilies with the highest contribution (Supplemental Fig 5B, Methods). We then built a heatmap providing a comprehensive overview of TE enrichment for these 164 subfamilies across 4614 samples and six histone marks (Fig 2C). We observed that TE subfamilies within a family tend to feature similar enrichments, for instance, the enrichment in H3K9me3 appears to be generalised across many L1 subfamilies. The same is true for most family clusters. TE families also seem to have distinct histone mark preferences. For instance, within the top 20 most enriched subfamilies, H3K4me1, H3K27ac & H3K27me3 share a similar profile of MIR and L2 subfamilies while H3K36me3 features Alu and H3K9me3 L1 andd ERVL-MaLR (Supplemental Fig 5C).

This perspective further supports the antagonism between activating and repressing mark TE enrichments and highlighted several enrichment clusters (Fig 2C). The enrichment of L1 in H3K9me3 while it is depleted everywhere else may be linked to previously reported TE repressing mechanisms^24,25^. A similarly high association can be found with ERV elements, another family noted for being repressed^26–28^. The enrichment of Alu in H3K36me3 stands out and is further investigated in the following sections. We also notice that MIR is highly enriched in marks that are depleted for the other TE families (H3K4me1, H3K27ac, H3K4me3), which may be due to their association with enhancers^15^. We found a widespread enrichment of ERVs across the different histones albeit at lower levels and less consistently. Since ERVs subfamilies are small and might not be the most enriched based on our obs–exp metric, we also explored enrichment based on fold change (observed / expected). We found that ERVs, were the most enriched according to fold change while Alu, L1 and MIR elements were most enriched according obs-exp (Supplemental Fig 6).

Taken together, these results suggest that TE families seem to have distinct preferences for histone mark enrichment and shows the complex, yet conserved relationship between TE and the epigenome.

### The different TE families display different enrichment dynamics as they age

Since older TE instances are more likely to have been either degraded by mutations or co-opted by the host genome^29^, we expect TEs epigenetic profile to change based on their age. It was reported in Su et al.^7^, that older Alus had a profile more like enhancers in terms of epigenetic state, conservation and TF binding potential. Pehrsson et al.^12^ also showed that DNA methylation of Alu went down with age. To determine if the age relationship was something that could be generalised across all TE families, we performed linear regressions of TE enrichment (mean across cell types for each subfamily) as a function of TE age (Fig 3A-B, Supplemental Fig 7). We observed that within some families (L1, ERVL-MaLR), many of the younger subfamilies were depleted in some mark (H3K27me3, H3K36me3) while enriched in others (H3K9me3) (Fig 3A). At the same time, for L1 and ERVL-MaLR, we found a general pattern of TEs within a family approaching 0% enrichment as they age (0% enrichment being the equivalent of being near the random simulations). This is consistent with TEs being degraded into background as they age^30^, which appears to be true whether they show an enrichment when they are young (H3K9me3) or a depletion (H3K27me3, H3K36me3).

**Figure 3.**
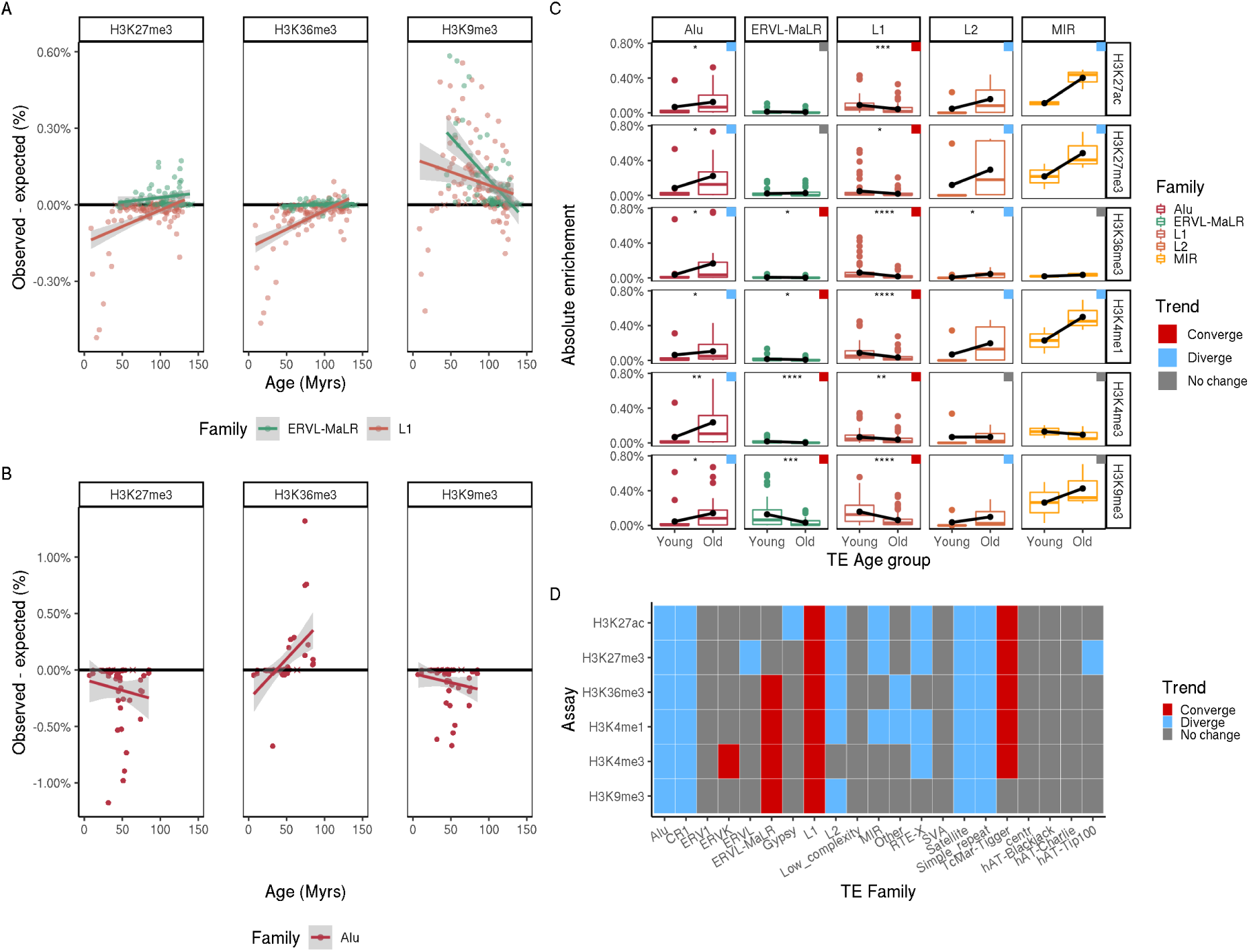
TE enrichment follows a family and context dependent age continuum. **A|** TE Family enrichment (obs-exp%) in function of estimated age for L1 and ERVL-MaLR in H3K27me3, H3K9me3 and H3K36me3. Each point is the mean enrichment (obs-exp%) of a TE subfamily across all samples. Black line is at 0% enrichment. Line shows linear regression fit, crosses are small sized subfamilies excluded from regression., **B|** Same as A for Alu TE family. **C|** Absolute enrichment (distance from 0 obs-exp% enrichment) of Young and Old TE subfamilies. Black line connects the means of age groups. (*: p <= 0.05,**: p <= 0.01,***: p <= 0.001,****: p <= 0.0001) Color in corner represents the trend. **D|** Dynamic of TE enrichment between young and old. Categorises the observations of **C** by dynamics and includes additional families. No change: no significant change, Diverge: Older TE have higher Absolute enrichment, Converge: Older TE have lower Absolute enrichment thus converge to 0.

While we found that some TE families tend towards no enrichment as they age, we also found a few families that diverged from the expected level. Alus show a distinct decline in enrichment in H3K4me3, H3K9me3 and H3K27me3 while for H3K36me3 it was an increase (Fig 3B, Supplemental Fig 7B). There are also many cases where there were little to no differences as they aged, such as for ERVK, ERVL and ERV1 (Supplemental Fig 7C). This exposes that the relationship between TE age and enrichment in the epigenome is complex and varies depending on family. We also looked at subfamily average length and instance count as potential cofounders and found that there were some correlations with TE age (Supplemental Fig 8). This means that those properties could also be associated with the observed TE enrichment evolution over time. In addition, since younger TEs tend to be harder to characterize due to mappability issues (less time for mutations to generate unique reads), we investigated the mappability and TE age correlation and only found it to be noticeable within L1 and Alu (Supplemental Fig 9A). However, we note that the less mappable TEs were not necessarily less enriched (Supplemental Fig 9 B,C) and thus, unlikely to be the cause of the observed enrichments.

Overall, two contrasting age dynamics were found: some TE families become more enriched or depleted as they age and others tend towards the expected background. To better characterise these dynamics, we measured the absolute enrichment of old and young TEs. We split the top 50% oldest and youngest TE subfamilies per group (Methods). We observed once again that some families had diverged from expectation as they aged (Alu, L2, MIR) while others converged to expectation (L1, ERVL-MaLR) (Fig 3C). These results are also summarized with more TE families in Fig 3D.

Taken together, these results highlight the complex TE enrichment association with histone marks that evolved differently depending on the TE family.

### TE enrichment varies between cell types and histone mark contexts

Next, we were interested in whether TEs could be involved in distinct cellular profiles and we measured the association of enrichment with cell types. Since the TE enrichment was primarily associated with the assay, we first grouped all samples per assay and then performed t-tests of cell type’s TE enrichment (significant only) against the mean for each histone mark and TE family pairs and sorted the cell types (Methods, Supplemental Table 1, 2).

For instance, looking at Alus within H3K36me3 due to their unexpected enrichment (Fig 2C), one notable cell type was the colon, which were significantly more enriched than the mean (10.9% average obs-exp versus 5.5% Fig 4A). We also observed high variability in some tissue’s enrichment. Indeed, looking at brain, lymphocytes of B lineage or endoderm derived structure, we found some samples enriched from among the lowest across all cell types to the highest (Fig 4A). Since each of our cell types were in fact heterogeneous groupings, we then investigated the underlying original cell labels. We found that the variability was due, at least in part, to the underlying cell label within the assigned cell types (Fig 4B). This exposed a high variability of TE enrichment across cell labels, even within a given cell type. We found that the Alu enrichment within lymphocytes of B lineage was mainly driven by the *B cells* (mean 19.3%) and not observed in tonsil germinal center B cell (mean 1.1%). Conversely, the colon cell type had more consistent results and smaller variance in its enrichment.

**Figure 4.**
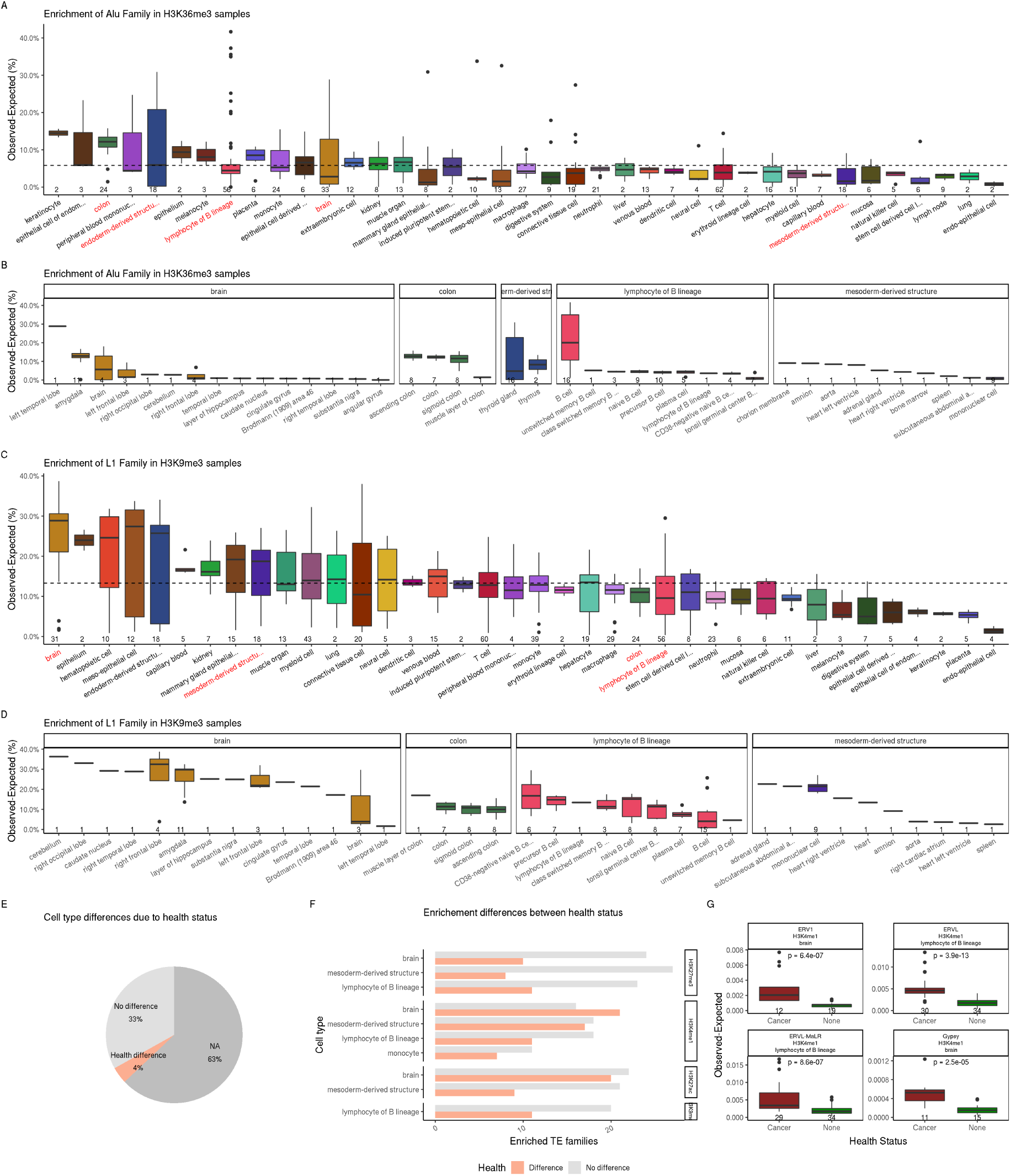
TE are enriched in histones in a cell type specific manner. **A|** Mean Alu Enrichment in H3K36me3 samples grouped by cell type. Each point is the mean of all Alu enrichment (obs-exp%, significantly enriched only) for one sample. Boxplot shows distribution across samples, number of samples for each cell type listed below. Sorted by category Mean enrichment. Dashed line, overall mean. Only cell types including more than one enriched sample are shown. Highlighted in red are Cell types displayed in B. **B**| Underlying cell type label within select cell types displayed in **A. C-D|** Same as figure A and B for L1 in H3K9me3 **E|** Proportion of cases (cell type-histone pairs and TE combinations) where there were significant differences depending on sample’s health status. **F|** Top 10 Cell type – Histone pairs with the most TE’s featuring significant differences between health statuses. X axis is the number of TE families featuring the significant difference (orange) or having no difference (gray). **G**| Four examples of TE enrichment (obs-exp, significantly enriched only) significant differences within cell type – Histone pairs.

We were also interested in the L1 enrichment observed in H3K9me3 (Fig 2C). There, brain had the highest enrichment (mean 24.1% versus 13.3% across cell types Fig 4C), which is line with reports of high brain and L1 associations in different contexts. L1s are upregulated in neural cell differentiation, in stress (rat hippocampus), brain diseases and for mosaicism while also being repressed by H3K9me3^31–34^. In contrast to Alu in H3K36me3, keratinocyte, colon and lymphocytes of B lineage had lower enrichment relative to mean (Fig 4A-C). This shows that the TE associations with histone marks change between tissues. Notably, within the brain cell type we find that differently annotated samples could have drastically different levels of enrichment (37% to 3%) further supporting that better and finer cell type annotation may help better understand the contribution of TEs (Fig 4D).

Finally, we tested if the health status of the samples had an impact on the enrichment levels observed in the different cell types. Of the cell types-histone pairs with multiple health status available (2024, 33% of samples), we found that 218 cases (4% of the total) of TEs featuring significant differences between health statuses (Fig 4E, Supplemental Fig 10). We highlight the top 10 of the cell types-histone pairs that had the most significant health associated differences (Fig 4F). For instance, we find that in H3K4me1 brain samples TE enrichment was significantly distinct between healthy and cancer cells across 21 TE families. We also show the top 4 cases with the most significant differences between healthy and cancer samples (Fig 4G). In all those cases, we find that cancer samples had a significantly higher TE enrichment.

These results show that different TEs are associated with different cell types and that health status can also affect those associations.

### Identifying notable TE candidates from cell type specific enrichments

Having found clear associations between TEs and cell types, we wanted to know how specific these associations were and if we could leverage their specificity to identify notable associations. To get an idea of the cell type specificity of TE enrichments, we first grouped all samples by cell type and histone mark and measured the proportion of samples in which each of the select TEs were significantly enriched (Supplemental Fig 11A). We found that most TEs were enriched in only a few cell types, except for H3K9me3 samples enriched across most cell types. Next, we added up the number of cell types in which TE subfamilies were enriched, to measure the cell type specificity of TE enrichments and confirmed that it differed based on histone marks (Fig 5A). H3K9me3 featured multiple families that were enriched across many (>30) cell types while H3K4me3 and H3K27ac are highly specific with few (<30) cell types enriched. In particular, L1 and ERV TEs were enriched across cell types in H3K9me3, suggesting these TEs are repressed in a non-cell type specific way. We observed that the MIR family had the opposite trend. Although a family with few subfamilies, the number of cell types in which these subfamilies were enriched was high (~43) in H3K27ac and low (~10) in H3K9me3.

**Figure 5.**
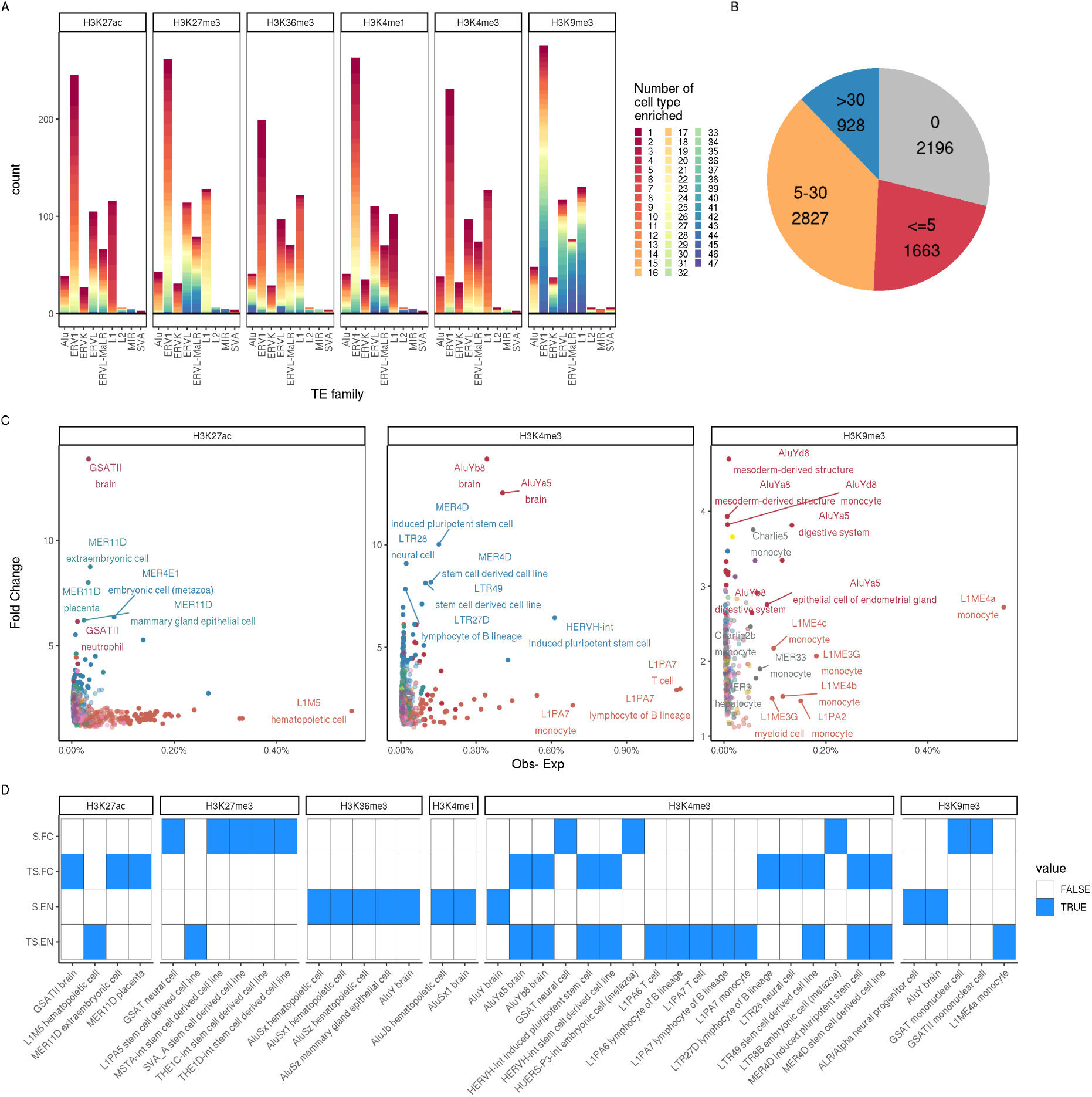
Identifying notable TE candidates from cell type specific enrichments. **A|** Number of TE subfamilies within each family that were enriched for each number of cell types. X axis shows the different TE families, Y axis shows the count of enriched TE subfamilies and the color shows the number of cell types the enrichment was observed in (Red=Specific, Blue=Non-Specific).**B|** Pie chart of the number of subfamilies enriched in 3 bins of cell type numbers (or not enriched, gray) across all histones **C|** Fold change and obs–exp % enrichment of significantly enriched TE subfamilies per cell type of cell type specific subfamilies (enriched in 5 or less cell types, red segment in B). Labels are a random subset of the candidates: most enriched (95^th^ percentile) points in terms of obs – exp or fold change. Only 3 histones are shown (H3K27ac, H3K4me3 and H3K9me3). **D|** Top 40 (10 selected per method) of the putative TE candidates annotated by the method of determination in y axis. S.FC is surplus fold change, TS.FC is top specific fold change, S.EN is surplus enrichment (obs-exp) and TS.EN is top specific enrichment (Obs-exp).

In summary, we found that across all marks, 1663 (21.8%) subfamilies were enriched in 1 to 5 cell types, 2827 (37.1%) in 6 to 30 cell types and 928 (12.2%) in 30 or more cell types (out of 47 ell types, Fig 5B). We catalogued the TE subfamilies that were enriched in a cell type specific manner (enriched in 1-5 cell types) and found that ERV1 subfamilies had the most specific TE enrichments across all marks (Supplemental Fig 11B). Broken down by histone mark, we find that for H3K9me3 only 151 (11.9%) were enriched in 1 to 5 cell types compared to 361 (28.4%) for H3K27ac (Supplemental Fig 11C).

Finally to identify TE to cell type associations for further analyses, we selected TE candidates through two approaches: (i) cell type specific TE association with high enrichment (top specific obs-exp (TS.EN) or fold change (TS.FC)) and (ii) TEs that were much more enriched for a cell type than all the others (surplus obs-exp (S.EN) or fold change (S.FC)). For the top specific enrichment, we selected as candidates the TE-Cell type-Histone triplets that were more enriched than the 95^th^ percentile of their histone group for either enrichment metric (Fig 5C and Supplemental Fig 12A). This led to the selection of 219 triplets based on obs-exp (TS.EN) and 219 triplets based on fold change (TS.FC, Supplemental Table 3). For instance, in H3K4me3, AluYb8 TE subfamily was selected since it had the highest fold change (14.19) in brain. Similarly, also in H3K4me4, L1PA7 was selected since it had the highest obs-exp enrichment (1.11%) in T-cell.

For the highest surplus, we identified the TEs that were much more enriched for a cell type than all the others by calculating the difference between the individual cell type TE enrichment and the mean across all cell types for each subfamily. We selected as candidate the top 20 TE-cell type pairs with the most difference in terms of obs-exp enrichment (S.EN, Supplemental Fig 11D) and Fold change (S.FC, Supplemental Fig 11E). The enrichment surplus was dominated by Alu subfamilies associated with Brain, hematopoietic cells and mammary gland epithelial cells. For instance, AluY was 5.6% enriched for H3K9me3 in the brain, 4.5% more than the average across cell types (1.1%). Meanwhile, fold change surplus was mostly associated with GSAT centr repeats and ERV TEs. THE1C-int in H3K27ac had a 16.5 fold change enrichment while the mean fold change was only 2.8 fold. From all of this, we compiled a list of 456 cell type-histone-TE candidate triplets with their methods of identification (438 from cell type specific TE associations and 40 from the surplus criteria, with some being shared) (Fig 5D, Supplemental Table 3). Among the candidates, we note multiple previously observed associations such as MER11D in placenta^35,36^, SVA_A for Stem cell being in line with pluripotent cell associations^11^and many Alus being enriched in brain^37,38^.

These observations highlight how our candidate selection managed to recapture many previous associations and suggests that some of these new candidates could serve as cell type specific regulatory elements and are worth further investigation.

### Identified TEs candidates are associated with relevant cell type biological processes and genes

Finally, we wanted to determine if we could link any of our 456 putative candidate TE subfamilies to potential activity through gene ontology associations. We merged the samples from a given candidate triplet to have one aggregate representative per context (merging of all peaks across samples from a cell type, assay and TE subfamily, Methods). For each merged sample we also generated a TE subfamily control keeping only the select TE’s instances that did not have peaks in the aforementioned representative sample (Supplemental Fig 13). This was to check the importance of overlapping the histone within cell type for our triplets. Next, we looked at the biological processes terms that were significant compared to genomic background with GREAT^39,40^ (Methods). For the 31 H3K27ac triplets, we found that there was some similarity in enriched terms between samples covering the same TEs, even across cell types (Fig 6A, Supplemental Table 4). Across marks, a similar pattern was observed for a subset of 209 triplets (Supplemental Fig 14, 15). It was clear that different processes were enriched within different triplets and thus, that TEs were, on some level, associated with different biological processes.

**Figure 6.**
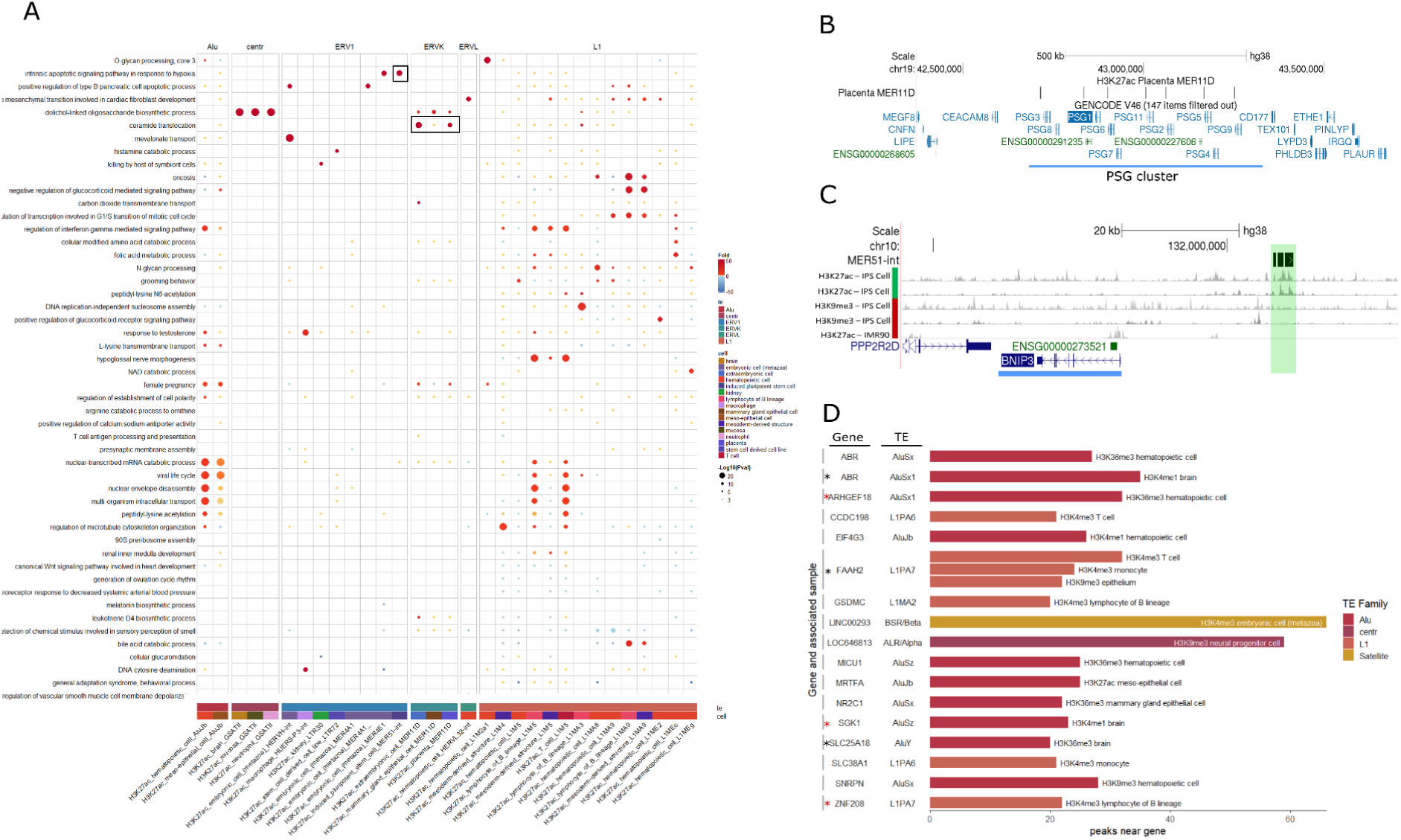
TEs associates with cell type relevant biological processes and genes. **A|**Fold change enrichment difference from TE control of GO biological processes within H3K27ac candidate triplet (TE-Assay-Cell Type) samples. (fold change data – fold change of associated TE control) red enriched, yellow between −1 and 1, blue depleted. Circle sizes represents significance of the enrichment. Terms selected based on data, favored most enriched terms per candidates. In rectangle are some mentioned processes. **B|** Genome tracks around *PSG* gene cluster (blue highlight) showing a cluster of MER11D overlapping peaks from H3K27ac placenta sample **C|** Genome tracks around *BNIP3* gene (blue highlight) showing a few nearby MER51-int instances (green highlight) and IPSC H3K27ac and H3K9me3 tracks, as well as an H3K27ac IMR90 sample track, all from the IHEC portal (part of the underlying samples, but distinct from the actual data used due to the reprocessing). **D|** number of peaks near genes (within 50kb) found within candidate triplets samples. Colored by TE family, only top gene per sample shown. star: supported RNA/protein expression data, red star: weaker support.

For example, we investigated MER11D in placenta and the associated biological processes and found enrichments for female pregnancy, ceramid and sphingolipids translocation (Supplemental Fig 14, 15A), all biological processes that are also involved in pregnancy and placenta^41,42^. We found a cluster of peaks within the PSG gene cluster (Fig 6B), near the *ABCB1* gene (and overlapping *RUNDC3B*) and near the *EPO* gene where a peak overlapped a candidate cis-regulatory elements (cCRE) (Supplemental Fig 16A, B). The *PSG* gene cluster stands for pregnancy-specific glycoprotein is directly related to pregnancy, while *ABCB1* is associated with ceramid and transports proteins and *EPO* promotes blood cells and is secreted from fetal liver.

The enrichment of oxygen associated terms in MER51-int H3K27ac iPSCs also stood out as they were highly significant and high in fold change (cellular response to hypoxia, 4 hit, 1005 fold enrichment, pval= 9.37E-12; response to hyperoxia, 6 hits, 67 fold enrichment, pval=4.92E-10; regulation of release of cytochrome c from mitochondria, 8 hits, 42 fold enrichment, pval=1.95E-11; Supplemental 14, 15B). With the triplet (MER51-int H3K27ac iPSCs) containing 59 peaks, these terms accounted for between 6.78% to 13.56% of the peaks in this cell type. This is consistent with the fact that oxygen levels are known to be important in iPSCs and pluripotency^43^. When we looked at the genes that explained these biological processes enrichments, we found 3, *BNIP3*, *HDAC2* and *HGF*. Among those, we highlight *BNIP3* a gene linked to the 3 terms and near (within 50kb) 5 MER51-int associated peaks in H3K27ac. These peaks were missing from other cell types and were not found in H3K9me3 samples (Fig 6C). This shows an example of a TE, MER51-int, being associated with activating histone mark H3K27ac near a gene associated with biological processes relevant to a cell type of interest. We also note that these peaks did not overlap previously annotated cCREs, even if some were close (Supplemental Fig 16C). We found that AluY H3K4me3 peaks in brain were associated with autophagosome and many peaks around two brain related genes, *SYT11* and *RIT1* (Supplemental Fig 15C, 16D). Finally, we looked at genes near peaks and found multiple cases (20-31%; Fig 6D, star annotated genes) of the gene or protein product being expressed in the candidate’s cell type. For instance, 32 peaks from L1PA7 H3K4me3 T-cell triplet were closest to *FAAH2* (Fig 6D) a gene highly expressed in T-cell according to The Human Protein Atlas (www.proteinatlas.org, Supplemental Fig 17A)^44^. We also found *SLC25A18* (closest to 20 peaks) within H3K36me3 AluY Brain triplet being mainly expressed in brain and *ABR* (closest to 35 peaks) for H3K4me1 AluSx1 Brain triplet was also most expressed in brain (Supplemental 17B, C).

Taken together, these results show that through our candidate selection, we could find cases of TE instances in proximity to genes associated with biological processes relevant to their cell type.

## Discussion

A number of recent studies have suggested a role for TEs in genome regulation ^10,31,45–49^. In this context, a comprehensive study of TE and their association with the epigenome was needed. Here we investigated, 4614 samples spanning 175 cell annotations from 47 cell types and 6 histone marks. On average 55.8% of peaks were found to overlap with TE and that the overlap varied greatly depending on the histone mark investigated. This is in line with the observations by Pehrsson et al.^12^ where they found varying degrees of TE overlap depending on epigenetic state. We also observed that H3K9me3, a less investigated repressive mark, had far more overlap with TEs (78.3%). We find that H3K9me3 is enriched especially in L1 and that this enrichment was not cell type specific. There was almost no enrichment in L1 for other histone marks. The role of H3K9me3 in silencing TE was established before^22,24,31^ and was particularly highlighted in the context of brain tissue^25,34^. Our findings suggest that L1 is specifically being targeted by H3K9me3 histone modification across tissues to silence it.

While observing an association of TEs with a repressing mark was expected, we also found distinct enrichment of Alus with H3K36me3. Alus are one of the few TE families still able to transpose in the human genome^51^. In contrast to L1, they featured little association with H3K9me3. We also found that ERVs TEs were the most widespread TEs with some level of enrichment across most histone marks further supporting the ERV’s contribution to the transcriptional landscape^47,52^.

The TE-age association is a perspective that is worthy to explore as it relates to the time a TE had to be co-opted or decay within the genome. It was previously found that Alu and SINE become less methylated as they age^12^, Alu become more preferred by H3K36me3 as they age^7^, that older SINE are more in open chromatin and that generally TEs lose their motifs as they age^53^. Su et al. proposed Alus as proto-enhancers due to their general properties becoming more enhancer like as they aged and we were interested in seeing if that observation held true for other TEs. We found that while their general observation for Alu held true, the overall relationship between other TEs and TE age was complex. We find that in L1 and ERV TE families, whether they start enriched or depleted, TEs tend toward no enrichment as they get older. This can be interpreted as most TE degrading into background which is supported by previous results^53^. Notably, we found that some TE families (Alu, MIR, L2, Fig 3D), instead diverged from no enrichment as they got older, indicating they either became more enriched or depleted. This can be interpreted as most TE degrading into background which is supported by previous results^53^. Our results highlight varying TE family associations with histone marks and evolutionary trajectories over time.

The idea that TEs are involved in a cell type specific way has been supported by many studies ^16,54^ and here, with our large and varied dataset, we aimed to test and establish to what extent the association of TEs and the epigenome was cell type specific. We found that TEs can be enriched within histone marks in a cell type specific manner. While most cell types have a similar TE enrichment, select tissues displayed significantly distinct enrichments depending on TE and histone mark. Alus in H3K36me3 were significantly more enriched in colon than most other tissues. We also found a high enrichment within Lymphocytes. The importance of TEs in the immunity has been reported before ^10,45,55,56^ and here we also observe high variability which is a factor of developing importance in the context of immunity^57^. L1 TEs were more enriched in brain than most other TEs within H3K9me3, which supports previous reports of L1 associations with brain^31,33,34^ and H3K9me3^24,28,58^. We also found that in 4% of cell types-histone combinations cancer (or disease) samples could also feature significantly different TE enrichments than healthy ones. However, due to our datasets limited focus on health status contrast, we may be missing a lot of health associated TE enrichment differences.

We also measured cell type specificity of TE enrichments, through the number of cell types in which they were enriched. We found that TEs tended to be specific for most subfamilies in the activating marks while they were more non-specific subfamily in H3K9me3 (Fig 5A, supplemental Fig 13C). In contrast, we note that MIR elements were non-specific in H3K27ac while specific in H3K9me3 (Fig 5A). Given previous reports of MIR enhancer activity^15^, we speculate that MIR is widely present in enhancers but selectively repressed for cell type specific purposes.

Finally, we used a set of criteria to identify 456 enriched cell type-histone-TE candidates (Fig 5D,G). We found that within our candidates, some biological processes were enriched in specific triplets. Among our candidates we identify MER11D in H3K27ac related to placenta and extraembryonic cells. The association between MER11 and placenta was observed before^35,36^ and we found a cluster of MER11D associated peaks within the *PSG* gene cluster. We also highlight a MER51-int H3K27ac iPSCs enrichment in oxygen associated terms (Fig 6A). This enrichment came in part from MER51-int elements near *BNIP3* gene which were missing from repressive mark and some other cell types (Fig 6C). Looking directly at genes, we found that the genes closest to our peaks, tended to have been reported as expressed within the triplet’s cell type. However, the fact that the associations came from peak clusters near the same genes might be a confounding factor.

While our large and heterogeneous dataset enabled our comprehensive study, it also came with some limitations. First although a harmonized reprocessing was made through a singular pipeline for all samples, some batch effect from the different consortium could still be detected. However, since different consortium mostly investigated different cell types, the specific cell types within the categories would also influence such differences. Additionally, we observed many cell type specific enrichments but it would be interesting to perform a similar TE analysis on single cell data to better capture the differences between cell types which could be lost in our bulk and aggregated data. Finally, due to the ambiguity that comes from multi-mapped reads, only uniquely mapped reads were used for in this study. Since TEs can lead to multi-mapped reads^59^, it is likely that some TE reads, and thus TE peaks, may have been lost leading to underestimating enrichments. It would be interesting to assess the exact TE contents lost from multi-map reads in future studies.

In summary, our data present an comprehensive overview of TE contents across histone marks and cell types. It shows the consistent yet complex relationship between TEs and the epigenome and further supports the implication of TEs in genome regulation.

## Supporting information

Supplemental figures and table desc

## Acknowledgements

This work was supported by a Canadian Institutes of Health Research (CIHR) program grant (CEE-151618) for the McGill Epigenomics Mapping Center, which is part of the Canadian Epigenetics, Environment and Health Research Consortium (CEEHRC) Network. G.B. is supported by a Canada Research Chair Tier 1 award and a FRQ-S, Distinguished Research Scholar award. We would like to acknowledge Calcul Quebec and the Digital Research Alliance of Canada for access to computing resources. We thank IHEC for making available reprocessed and harmonized epigenomic data from a large collection of human cell types^18^.

## Methods

### Data Collection

The dataset was downloaded from the IHEC EpiATLAS integrative analysis sFTP server on January 23^rd^ 2023 and also available on the IHEC data portal (https://epigenomesportal.ca/ihec/, https://ihec-epigenomes.org/epiatlas/data/). The available ChIPSeq narrowPeak files for the 6 main histone marks (H3K4me1, H3K4me3, H3K27ac, H3K27me3, H3K9me3, H3K36me3) were downloaded. The narrowPeaks were obtained using GRCh38 reference build, with pval of 0.01. The data downloaded included samples from 8 consortiums (Blueprint, CEEHRC, ENCODE, DEEP, NIH Roadmap Epigenomics, AMED-CREST, GIS, EpiHK). The cell type and all metadata annotations was taken from the IHEC metadata harmonization v1.2. Only samples with an Epirr id within the metadata v1.2 were used (ENCODE samples were dismissed). Comparison were made with Roadmap complete Epigenomes while only considering the 6 main histone marks. ENCODE dataset sample count for comparison was obtained by filtering within the Experiment matrix, epigenome dataset for the 6 main histones totalling 1426 available samples (https://www.encodeproject.org/, Jan 2024).

### Quality control and sample selection

For each sample, we measured the distribution of peaks within regions relative to transcription start sites (TSS). Peaks were attributed to the first group they fell within among: TSS (within 1000 bp from TSS), Promoter (within 5000 bp upstream of TSS), Intragenic (Overlapping genes), Proximal (within 10 kbp from TSS), Distal (within 100 kbp from TSS) and desert (further than 100 kbp from TSS). The coverage of each region was compared between samples from the same histone mark and samples that were an outlier (according to boxplots, above (75% quantile) + 1.5 * interquartile range (IQR) or bellow (25% quantile) - 1.5 * IQR) in the covered percentage for 2 different regions were discarded. In addition, samples that were outliers in terms of peak count when grouped by histone marks were also discarded.

### Dimension reduction on ChIP-seq samples

To have broad overview of our samples similarity and do dimension reduction to display the data, we counted the number of peaks within 10 kbp windows across the entire genome using bedtools^60^ intersect for each sample. We sorted the windows by variance across the samples and kept the top 21,000 windows before discarding the top 1000 to protect against potential irregularities (e.g. regions with very high coverage). This resulted in 20,000 windows upon which UMAP was performed with r umap package using default arguments with 400 epochs.

### Measuring TE enrichment

Histone mark peaks were annotated using the UCSC RepeatMasker track^6^. We resized the peaks to 200bp around their center and counted the number of times ChIP-seq peaks overlapped TE instances with bedtool intersect, when more than one TE overlapped the peak, the largest overlap was kept. As done before ^47,61^, to have a random baseline to compare against, we simulated for each sample a library of 200 bp random regions, with the same distribution of distances to nearest genes. This was done using the distribution of peaks described in Quality control and sample selection method. For each sample, the simulation was repeated 1000 times and we counted the incidence of observed count being higher than the random baseline for each repeat subfamily. A repeat subfamily was identified as significantly over-represented (enriched) when the over-represented incidence was greater than 995/1000 (p < 0.005). coverage percentages were measured as *peaks overlapping TE/sample (or simulated sample) peak count*. Observed – Expected metric was calculated by subtracting the expected coverage (resulting from mean across the 1000 simulations) from the observed coverage from the sample. Significant Observed – Expected was calculated using the only the repeat subfamilies identified as significantly overepresented (thus always positive, because observed in inherently higher than expected for each subfamily used).

### Selection of the most enriched TE subfamilies

The most enriched TE were selected by doing a cumulative sum of the positive (>0) enrichment (observed-expected) TE subfamilies (excluding the simple_repeats) across all samples. Upon this cumulative measure, the threshold was set as the value of the upper whisker of a boxplot, the value beyond which values are considered outliers. It was defined as *(75% quantile) + 1.5 * IQR*.

### Correlation between samples and TEs

The correlation between samples was done with a matrix keeping the count of peaks found within the 20,000 10 kbp windows with highest variance (as described in the Dimension reduction on ChIP-seq samples section above) on which we grouped samples by assay and calculated the mean peak count per assay for each window. Then, the correlation between the assays was calculated with the Corr function in R.

The correlation between TEs was done with a matrix keeping the Observed – Expected (%) enrichment of the selected TE families for each sample on which we grouped the samples by assay and calculated the mean enrichment per assay for each TE family. Then the correlation between the assays was calculated with Corr function in R.

### Estimation of TE Mappability, age and age categorization

TE mappability was calculated using the 50 bp Mappability track^23^ from the UCSC Genome Browser, which is a conservative estimate of the true mappability since most of the reads in IHEC were targeting 75 bp and mappability increases with read length. The coverage of all TEs (also UCSC track) by the 50bp Unique Mappability (Umap 50) track gave us the proportion of each TEs that could be uniquely mapped which we used as mappability metric. The age estimates of each TE were based on the sequence divergence (milliDiv value from RepeatMasker) as described in Bogdan et al.^45^ We first divided the milliDiv value of each TE by 1000 and then by 2.2×10^−9^, the substitution rate of the human genome to calculate the age. The age of each family and subfamily was then obtained from the mean of all their instance’s age. We categorized the subfamilies into Old or Young depending on the subfamily’s age rank within the family. The youngest 50% were categorised young and oldest 50% old. For the dynamic groupings, if the difference between the old and young TE absolute enrichment (absolute value enrichment) was less than the young absolute enrichment (did not double or halve) and the p-value was larger than 0.05, the context (TE family for Histone mark) was deemed to have no change. If there was change, when the Old TEs absolute enrichment was larger than the Young TE’s the context was categorized as diverging (moving away from 0 enrichment) and otherwise converging (approaching 0 enrichment).

### Candidate selection

We selected TE candidates through two approaches: (i) cell type specific TE association with high enrichment (top specific obs-exp (TS.EN) or fold change (TS.FC)) and (ii) TEs that were much more enriched for a cell type than all the others (surplus obs-exp (S.EN) or fold change (S.FC)). For the top specific enrichment, we selected as candidates the TE-Cell type-Histone triplets that were more enriched than the 95^th^ percentile enrichment of their histone group for either enrichment metric (obs-expected or fold change). For the surplus method, we calculated the mean TE enrichment across all cell types and subtracted it from each cell type’s enrichment. We selected the top 20 (for both obs-expected and fold change)

This resulted in a final set of 456 candidates. For a more restrained and manageable set, we selected a subset of the top 15 most enriched candidates per histone for top specific obs-exp and fold change using their respective metric (obs-exp and fold change, respectively). We thus had 90 top specific obs-exp candidates, 90 top specific fold change and the 40 surplus candidates. This resulted in a reduced set of 209 candidates (due to some candidates being identified by more than one method).

### Identification of associated GO terms

To identify the GO terms associated with TEs depending on histone or cell type, we first merged all bed files by cell type and histone to have 1 representative per doublet (e.g. peaks from H3K9me3 in Brain cells). We then split the merged samples by TE to have the peaks associated with each TE for all doublets and resulting in triplets (e.g. peaks from H3K9me3 in brain cells overlapping L1PA4). As a control to observe the influence of TEs on their own, triplet controls were made by taking all instances of the given TE (straight from repeatmasker) and removing all those that overlapped peaks from the given triplet. These triplet controls represent the instances of the TE not within peaks of the cell type and histone (ex: L1PA4 peaks that are not in H3K9me3 brain peaks). We obtained the associated GO terms from these Triplets and associated controls using the R version of GREAT (rGREAT)^39^. We used the default configurations (5kb upstream, 1kb downstream, up to 1000kb) and background (genomic background) for the analysis. To account for the use of genomic background, the same analysis was made on the controls which were used to assess if the enrichment observed could be explained by our controls (TE itself or Histone-Cell type doublets). For visualisation purposes, GO term p-value was capped at 10E-200. The GO terms to show are selected by sorting the terms by p-value first and fold-change second. The genes supporting the GO term associations as well as the location of relevant peaks were obtained by using the GREAT website (https://great.stanford.edu/great/public/html/index.php) version 4.04, hg38 and default configuration.

### Visualisation

Figure generation was done using R^62^, heatmaps were made using complexheatmap^63^

## References

1. Finnegan, D. J. Eukaryotic transposable elements and genome evolution. Trends in Genetics 5, 103–107 (1989).

2. Wells, J. N. & Feschotte, C. A Field Guide to Eukaryotic Transposable Elements. Annu Rev Genet 54, 539–561 (2020).

3. Ye, M. et al. Specific subfamilies of transposable elements contribute to different domains of T lymphocyte enhancers. Proceedings of the National Academy of Sciences 117, 7905– 7916 (2020).

4. Carey, K. M., Patterson, G. & Wheeler, T. J. Transposable element subfamily annotation has a reproducibility problem. Mobile DNA 12, 4 (2021).

5. Smit, A. F. A., Tóth, G., Riggs, A. D. & Jurka, J. Ancestral, Mammalian-wide Subfamilies of LINE-1 Repetitive Sequences. Journal of Molecular Biology 246, 401–417 (1995).

6. Smit, A. F. A., Hubley, R. & Green, P. RepeatMasker Open-4.0. (2013).

7. Su, M., Han, D., Boyd-Kirkup, J., Yu, X. & Han, J.-D. J. Evolution of Alu Elements toward Enhancers. Cell Reports 7, 376–385 (2014).

8. Kunarso, G. et al. Transposable elements have rewired the core regulatory network of human embryonic stem cells. Nat Genet 42, 631–634 (2010).

9. Ohnuki, M. et al. Dynamic regulation of human endogenous retroviruses mediates factor-induced reprogramming and differentiation potential. Proc. Natl. Acad. Sci. U.S.A. 111, 12426–12431 (2014).

10. Chuong, E. B., Elde, N. C. & Feschotte, C. Regulatory evolution of innate immunity through co-option of endogenous retroviruses. Science (New York, N.Y.) 351, 1083 (2016).

11. Barnada, S. M. et al. Genomic features underlie the co-option of SVA transposons as cis-regulatory elements in human pluripotent stem cells. PLOS Genetics 18, e1010225 (2022).

12. Pehrsson, E. C., Choudhary, M. N. K., Sundaram, V. & Wang, T. The epigenomic landscape of transposable elements across normal human development and anatomy. Nat Commun 10, 1–16 (2019).

13. Feliciello, I., Sermek, A., Pezer, Ž., Matulić, M. & Ugarković, Đ. Heat Stress Affects H3K9me3 Level at Human Alpha Satellite DNA Repeats. Genes 11, 663 (2020).

14. Mosch, K., Franz, H., Soeroes, S., Singh, P. B. & Fischle, W. HP1 Recruits Activity-Dependent Neuroprotective Protein to H3K9me3 Marked Pericentromeric Heterochromatin for Silencing of Major Satellite Repeats. PLOS ONE 6, e15894 (2011).

15. Jjingo, D. et al. Mammalian-wide interspersed repeat (MIR)-derived enhancers and the regulation of human gene expression. Mobile DNA 5, 14 (2014).

16. Cao, Y. et al. Widespread roles of enhancer-like transposable elements in cell identity and long-range genomic interactions. Genome Res 29, 40–52 (2019).

17. Bujold, D. et al. The International Human Epigenome Consortium Data Portal. cels 3, 496–499.e2 (2016).

18. International Human Epigenome Consortium, EpiATLAS - a reference for human epigenomic research. *In preperation*.

19. Roadmap Epigenomics Consortium et al. Integrative analysis of 111 reference human epigenomes. Nature 518, 317–330 (2015).

20. Moore, J. E. et al. Expanded encyclopaedias of DNA elements in the human and mouse genomes. Nature 583, 699–710 (2020).

21. Allshire, R. C. & Madhani, H. D. Ten principles of heterochromatin formation and function. Nat Rev Mol Cell Biol 19, 229–244 (2018).

22. Kabi, M. & Filion, G. J. Heterochromatin: did H3K9 methylation evolve to tame transposons? Genome Biology 22, 325 (2021).

23. Karimzadeh, M., Ernst, C., Kundaje, A. & Hoffman, M. M. Umap and Bismap: quantifying genome and methylome mappability. Nucleic Acids Research 46, e120 (2018).

24. Seczynska, M., Bloor, S., Cuesta, S. M. & Lehner, P. J. Genome surveillance by HUSH-mediated silencing of intronless mobile elements. Nature 1–9 (2021) doi:10.1038/s41586-021-04228-1.

25. Erwin, J. A., Marchetto, M. C. & Gage, F. H. Mobile DNA elements in the generation of diversity and complexity in the brain. Nature Reviews Neuroscience 15, 497–506 (2014).

26. Collins, P. L., Kyle, K. E., Egawa, T., Shinkai, Y. & Oltz, E. M. The histone methyltransferase SETDB1 represses endogenous and exogenous retroviruses in B lymphocytes. Proceedings of the National Academy of Sciences 112, 8367–8372 (2015).

27. Wang, Z. et al. Dominant role of DNA methylation over H3K9me3 for IAP silencing in endoderm. Nat Commun 13, 5447 (2022).

28. Maksakova, I. A. et al. H3K9me3-binding proteins are dispensable for SETDB1/H3K9me3-dependent retroviral silencing. Epigenetics & Chromatin 4, 12 (2011).

29. Chuong, E. B., Elde, N. C. & Feschotte, C. Regulatory activities of transposable elements: from conflicts to benefits. Nat Rev Genet 18, 71–86 (2017).

30. Blumenstiel, J. P. Birth, School, Work, Death, and Resurrection: The Life Stages and Dynamics of Transposable Element Proliferation. Genes 10, 336 (2019).

31. Ahmadi, A., De Toma, I., Vilor-Tejedor, N., Eftekhariyan Ghamsari, M. R. & Sadeghi, I. Transposable elements in brain health and disease. Ageing Research Reviews 64, 101153 (2020).

32. Hunter, R. G., McEwen, B. S. & Pfaff, D. W. Environmental stress and transposon transcription in the mammalian brain. Mobile Genetic Elements 3, e24555 (2013).

33. Erwin, J. A. et al. L1-associated genomic regions are deleted in somatic cells of the healthy human brain. Nat Neurosci 19, 1583–1591 (2016).

34. Muotri, A. R. et al. Somatic mosaicism in neuronal precursor cells mediated by L1 retrotransposition. Nature 435, 903–910 (2005).

35. Bi, S., Gavrilova, O., Gong, D.-W., Mason, M. M. & Reitman, M. Identification of a Placental Enhancer for the Human Leptin Gene *. Journal of Biological Chemistry 272, 30583–30588 (1997).

36. Frost, J. M. et al. Regulation of human trophoblast gene expression by endogenous retroviruses. 2022.04.26.489485 Preprint at 10.1101/2022.04.26.489485 (2022).

37. Jacob-Hirsch, J. et al. Whole-genome sequencing reveals principles of brain retrotransposition in neurodevelopmental disorders. Cell Res 28, 187–203 (2018).

38. Mehler, M. F. & Mattick, J. S. Noncoding RNAs and RNA Editing in Brain Development, Functional Diversification, and Neurological Disease. Physiological Reviews 87, 799–823 (2007).

39. Gu, Z. & Hübschmann, D. rGREAT: an R/bioconductor package for functional enrichment on genomic regions. Bioinformatics 39, btac745 (2023).

40. McLean, C. Y. et al. GREAT improves functional interpretation of cis-regulatory regions. Nat Biotechnol 28, 495–501 (2010).

41. Fakhr, Y., Brindley, D. N. & Hemmings, D. G. Physiological and pathological functions of sphingolipids in pregnancy. Cellular Signalling 85, 110041 (2021).

42. Enthoven, L. F. et al. Effects of Pregnancy on Plasma Sphingolipids Using a Metabolomic and Quantitative Analysis Approach. Metabolites 13, 1026 (2023).

43. Nit, K., Tyszka-Czochara, M. & Bobis-Wozowicz, S. Oxygen as a Master Regulator of Human Pluripotent Stem Cell Function and Metabolism. Journal of Personalized Medicine 11, 905 (2021).

44. Uhlén, M. et al. Tissue-based map of the human proteome. Science 347, 1260419 (2015).

45. Bogdan, L., Barreiro, L. & Bourque, G. Transposable elements have contributed human regulatory regions that are activated upon bacterial infection. 9.

46. Kojima, S. et al. Mobile elements in human population-specific genome and phenotype divergence. 2022.03.25.485726 Preprint at 10.1101/2022.03.25.485726 (2022).

47. Jacques, P.-É., Jeyakani, J. & Bourque, G. The Majority of Primate-Specific Regulatory Sequences Are Derived from Transposable Elements. PLoS Genet 9, e1003504 (2013).

48. Barnada, S. M., et al. *Genomic Features Underlie the Co-Option of SVA Transposons as Cis-Regulatory Elements in Human Pluripotent Stem Cells*. 2022.01.10.475682 https://www.biorxiv.org/content/10.1101/2022.01.10.475682v1 (2022) doi:10.1101/2022.01.10.475682.

49. Ardeljan, D., Taylor, M. S., Ting, D. T. & Burns, K. H. The Human Long Interspersed Element-1 Retrotransposon: An Emerging Biomarker of Neoplasia. Clinical Chemistry 63, 816–822 (2017).

50. Saksouk, N., Simboeck, E. & Déjardin, J. Constitutive heterochromatin formation and transcription in mammals. Epigenetics and Chromatin 8, (2015).

51. Mills, R. E., Bennett, E. A., Iskow, R. C. & Devine, S. E. Which transposable elements are active in the human genome? Trends in Genetics 23, 183–191 (2007).

52. Cowley, M. & Oakey, R. J. Transposable Elements Re-Wire and Fine-Tune the Transcriptome. PLOS Genetics 9, e1003234 (2013).

53. Du, A. Y., Chobirko, J. D., Zhuo, X., Feschotte, C. & Wang, T. Regulatory Transposable Elements in the Encyclopedia of DNA Elements. 2023.09.05.556380 Preprint at 10.1101/2023.09.05.556380 (2023).

54. Bogu, G. K., Reverter, F., Marti-Renom, M. A., Snyder, M. P. & Guigó, R. Atlas of transcriptionally active transposable elements in human adult tissues. bioRxiv 714212 (2019) doi:10.1101/714212.

55. Stenz, L. The L1-dependant and Pol III transcribed Alu retrotransposon, from its discovery to innate immunity. Mol Biol Rep 48, 2775–2789 (2021).

56. Kong, Y. et al. Transposable element expression in tumors is associated with immune infiltration and increased antigenicity. Nature Communications 10, 5228 (2019).

57. Chen, X. et al. Transposable elements are associated with the variable response to influenza infection. 2022.05.10.491101 Preprint at 10.1101/2022.05.10.491101 (2022).

58. Pezic, D., Manakov, S. A., Sachidanandam, R. & Aravin, A. A. piRNA pathway targets active LINE1 elements to establish the repressive H3K9me3 mark in germ cells. Genes Dev. 28, 1410–1428 (2014).

59. Goerner-Potvin, P. & Bourque, G. Computational tools to unmask transposable elements. Nature Reviews Genetics 19, 688–704 (2018).

60. Quinlan, A. R. & Hall, I. M. BEDTools: a flexible suite of utilities for comparing genomic features. Bioinformatics 26, 841–842 (2010).

61. Zhuang, Q. K.-W. et al. Sex Chromosomes and Sex Phenotype Contribute to Biased DNA Methylation in Mouse Liver. Cells 9, 1436 (2020).

62. R Core Team. R: A language and environment for statistical computing. R Foundation for Statistical Computing.

63. Gu, Z., Eils, R. & Schlesner, M. Complex heatmaps reveal patterns and correlations in multidimensional genomic data. Bioinformatics 32, 2847–2849 (2016).

