## Supplemental figures and table desc for "Transposable elements impact the human regulatory landscape through cell type specific epigenomic associations"

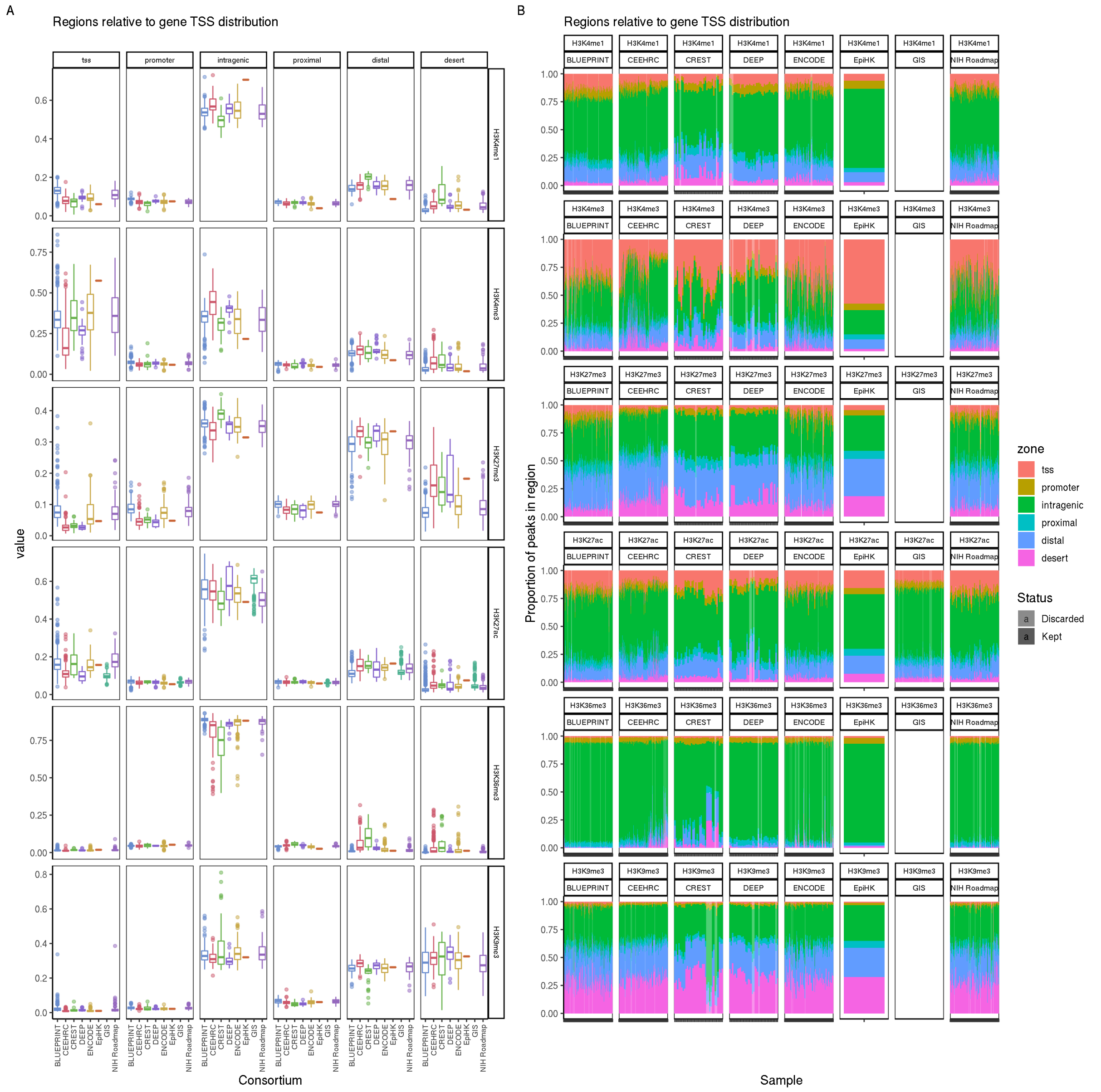


Supplemental Figure 1. Distribution of the peaks in regions relative to TSS.
**A)** Distribution of peaks within annotations relative to gene TSS. Horizontal facets are the different regions, the Y axis is the proportion found within the region **B)** X axis are the samples faceted by their assay and consortium. Y axis is the proportion found within the colored regions. Color are the different regions, transparent samples are those discarded due to outlier samples.

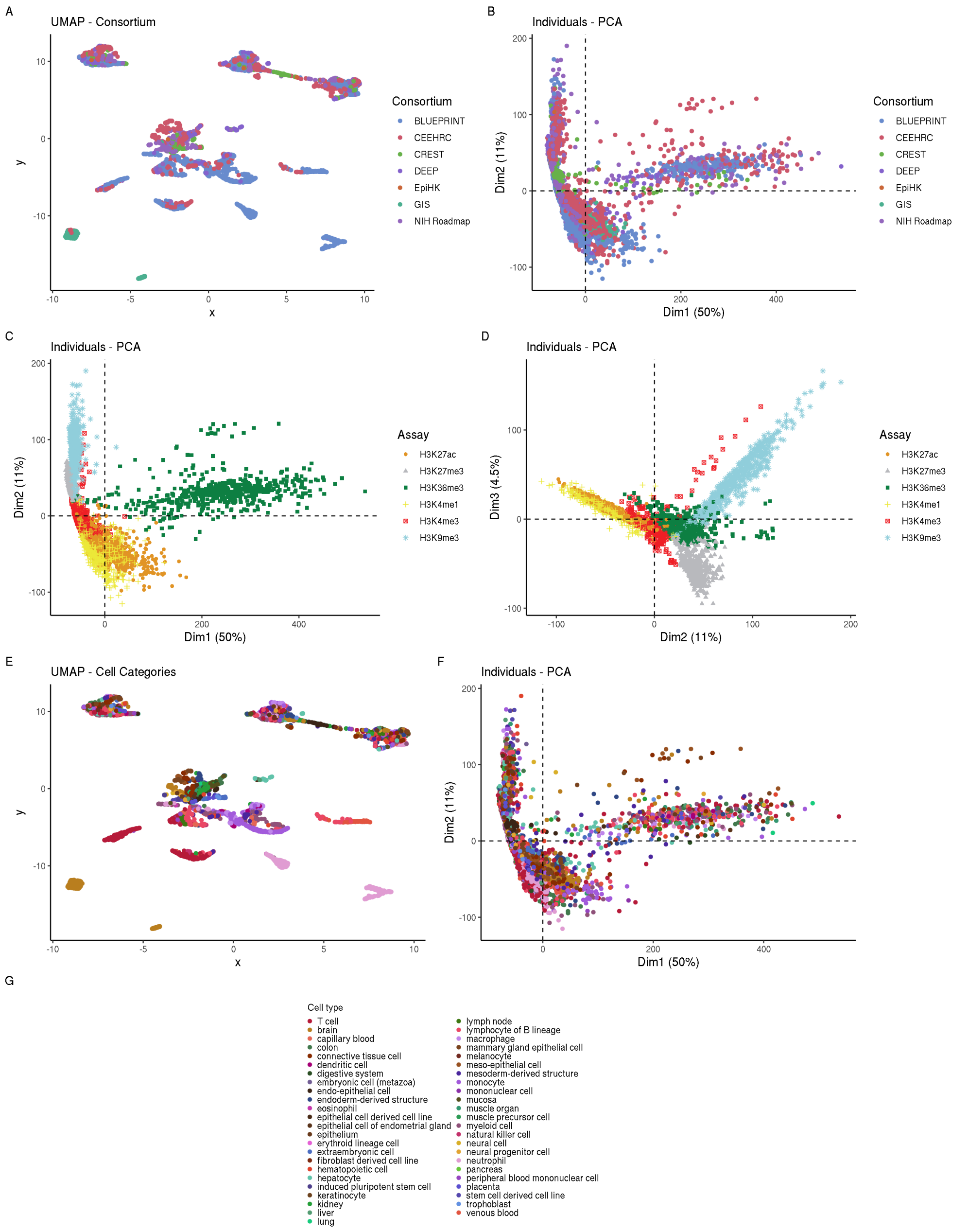


Supplemental Figure 2. ChIP-Seq samples data dimension reduction.

**A)** UMAP on peak count within 10kb regions across full genome of the 4614 samples. Only the 20,000 windows with most variance (excluding the first 1000) were used colored by consortium **B)** PCA of first 2 PCs using same data as A colored by Sample's Consortium **C)** colored by Sample's assay **D)** PCA of 2$^{nd}$ and 3$^{rd}$ PC colored by sample's assay **E)** Same as A colored by cell type **F)**PCA of first 2 PCs as in B and C colored by Cell type **G)** Cell type Color legend of E and F

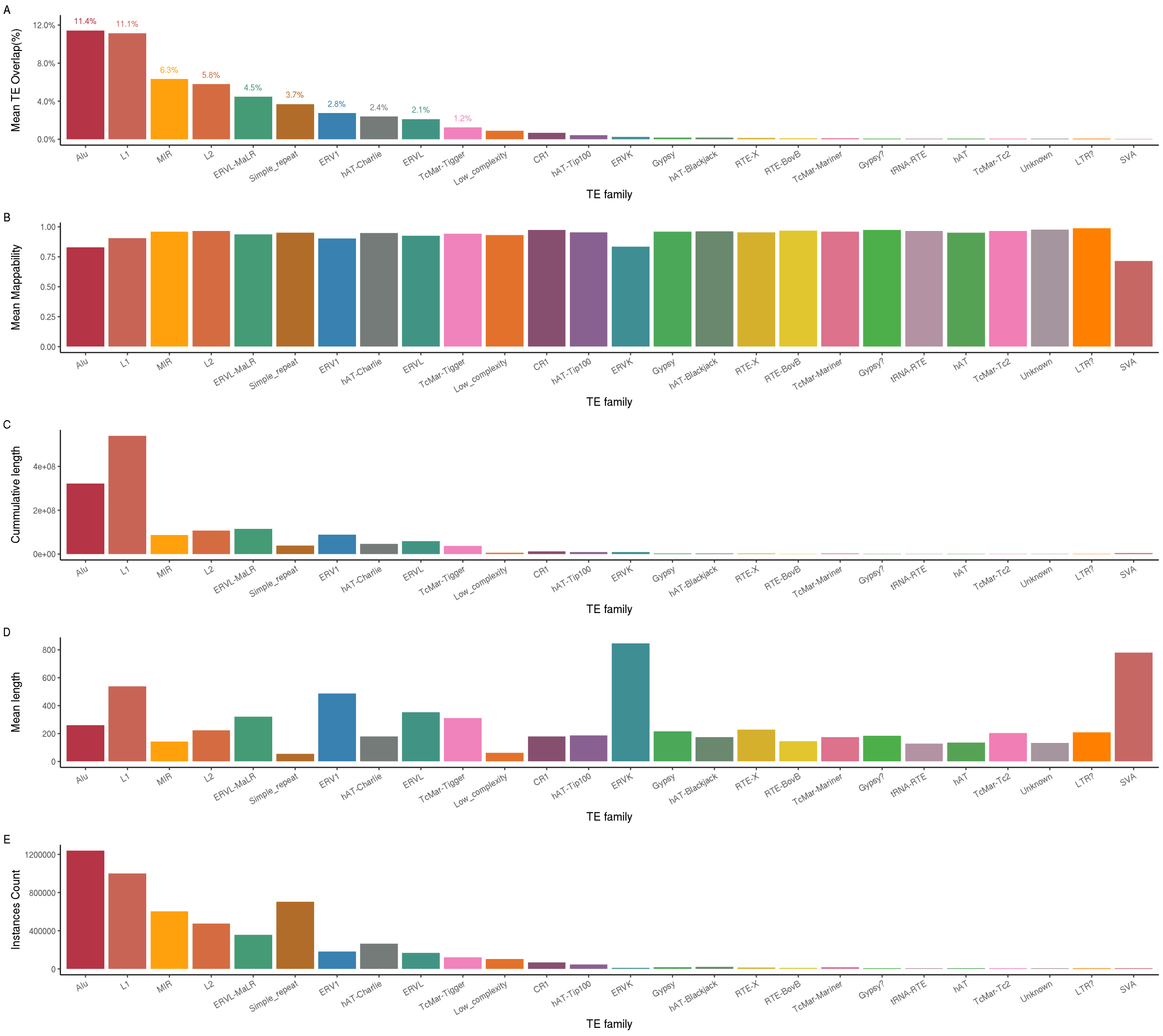


Supplemental Figure 3. TE Family Overlap and TE family properties.
**A)** Mean TE overlap across all samples **B)** TE Mappability estimates **C)** Cummulative bp length within the genome **D)** Mean TE instance length **E)** TE instance count. X axis sorted according to TE overlap(A). Only families containing at least 1000 instances shown. Centr family excluded due to extreme outlier mean length.

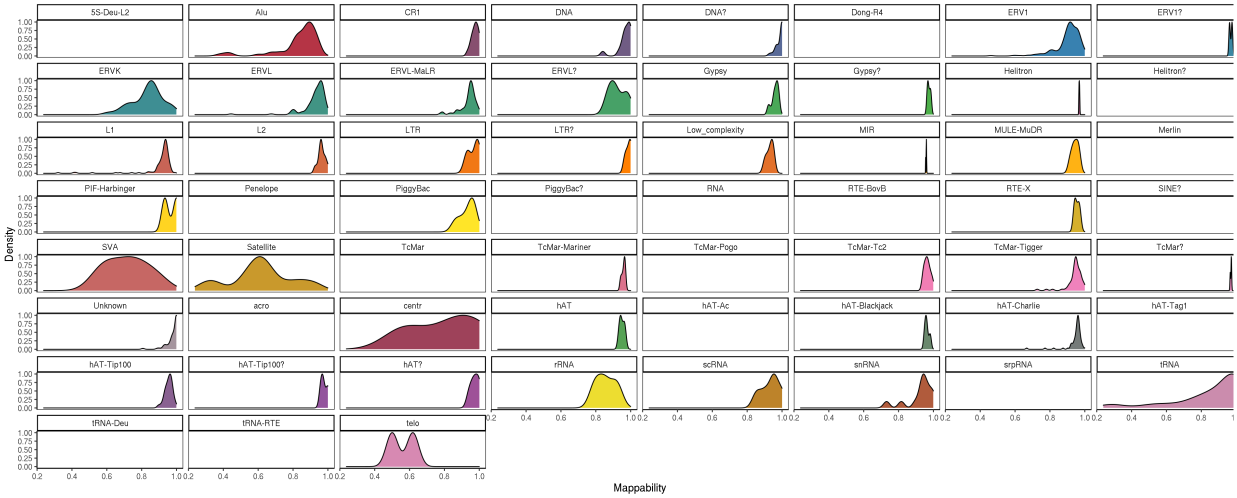


Supplemental Figure 4. TE Families Mappability distribution.
Density distribution of TE instances Mappability (TE coverage by the 50bp Unique Mappability (Umap 50) track) within the TE families. (Empty if not enough instances)

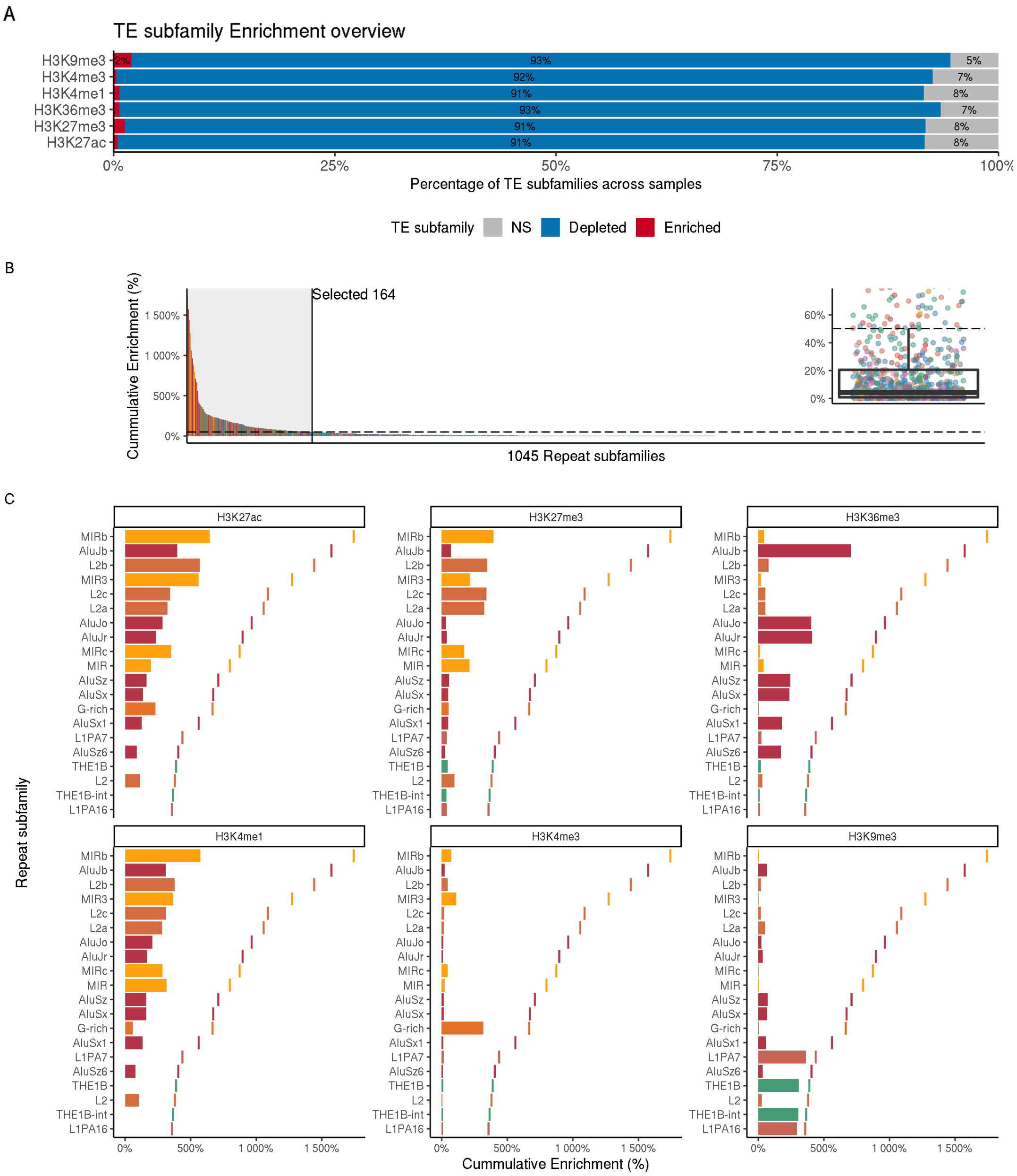


Supplemental Figure 5. Breakdown of TE Enrichments Within Subfamilies.
**A**) Percentage of TE subfamilies enriched or depleted relative to random simulation across assays. Each TE subfamily is enrichment is recorded individually for each sample. NS are non significantly enriched TEs **B)** Sorted cumulative sum of the 1045 TE subfamilies enrichment (obs-exp, significantly enriched, and therefore >0, only) across all samples. The 164 TE subfamilies above the threshold line (dashed line) were selected for the downstream analysis. The threshold was selected as the upper whisker from the boxplot of the cumulative sum of TE enrichment **C)** Top 20 of Cummulative sum of TE subfamilies genome overlap across all samples grouped by Histone mark. Vertical line is the sum of the enrichment across all assays. Thus, the bar size relative to the vertical line represents the proportion of the enrichment coming from the given assay.

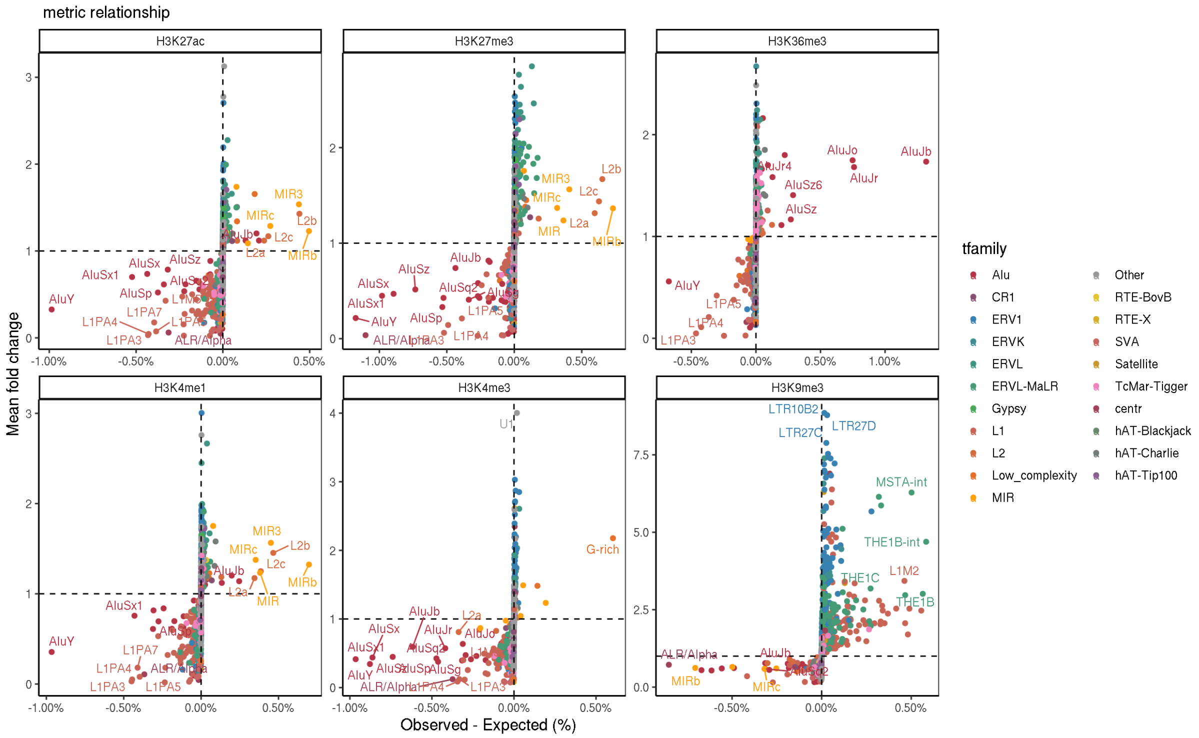


Supplemental Figure 6. Fold change Enrichment in function of Obs-Exp.
Mean Fold change in function of observed – expected enrichment of TE subfamilies (mean across cell types). Colored by TE family.

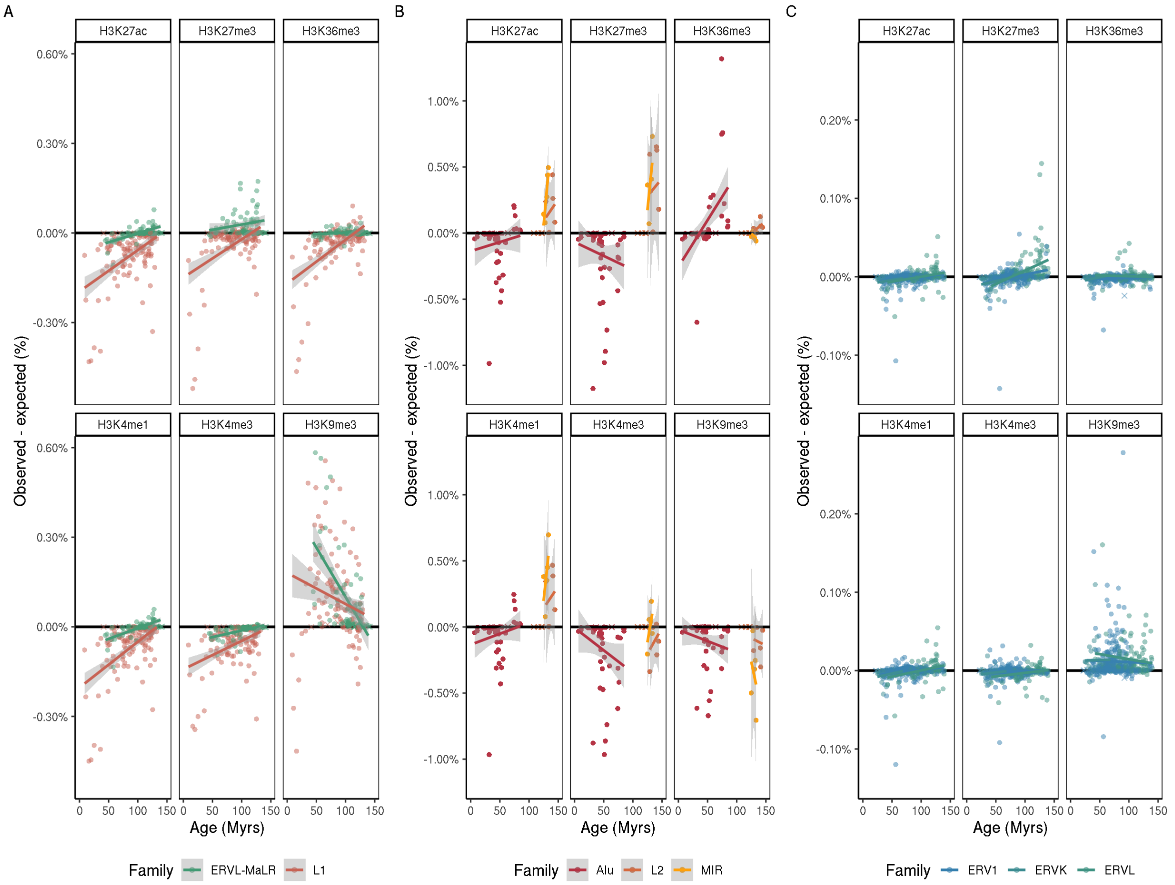


Supplemental Figure 7. Expanded age regression set.

**A)** TE Family enrichment (obs-expected%) in function of estimated age for L1, ERVL-MaLR across all 6 Histone marks. Black line is at 0 enrichment. Line shows linear regression fit, crosses are small sized subfamilies excluded from regression. **B)** For Alu,L2 and MIR **C)** For ERV1, ERVK and ERVL

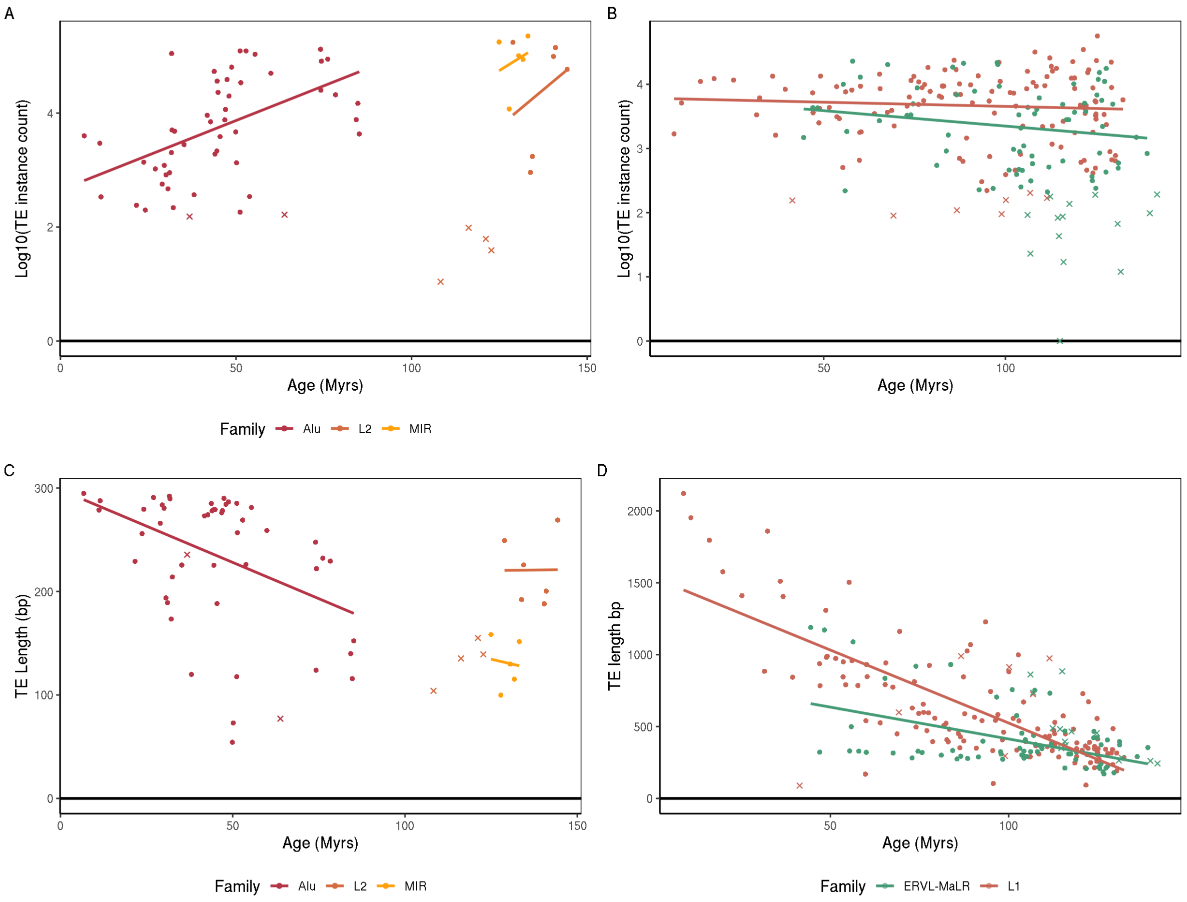


Supplemental Figure 8. Potential TE properties cofounding with age
**A)** Mean in instance count of TE subfamilies in function of their mean age estimate for Alu, MIR and L2. **B)** same as A for L1 and ERVL-MaLR. **C)** Mean TE Length of TE subfamilies in function of their mean age estimates for Alu, L2 and MIR. **D)** same as C for L1 and ERVL MaLR. X shapes are subfamilies too small (less than ) in instance count and were not used for regression.

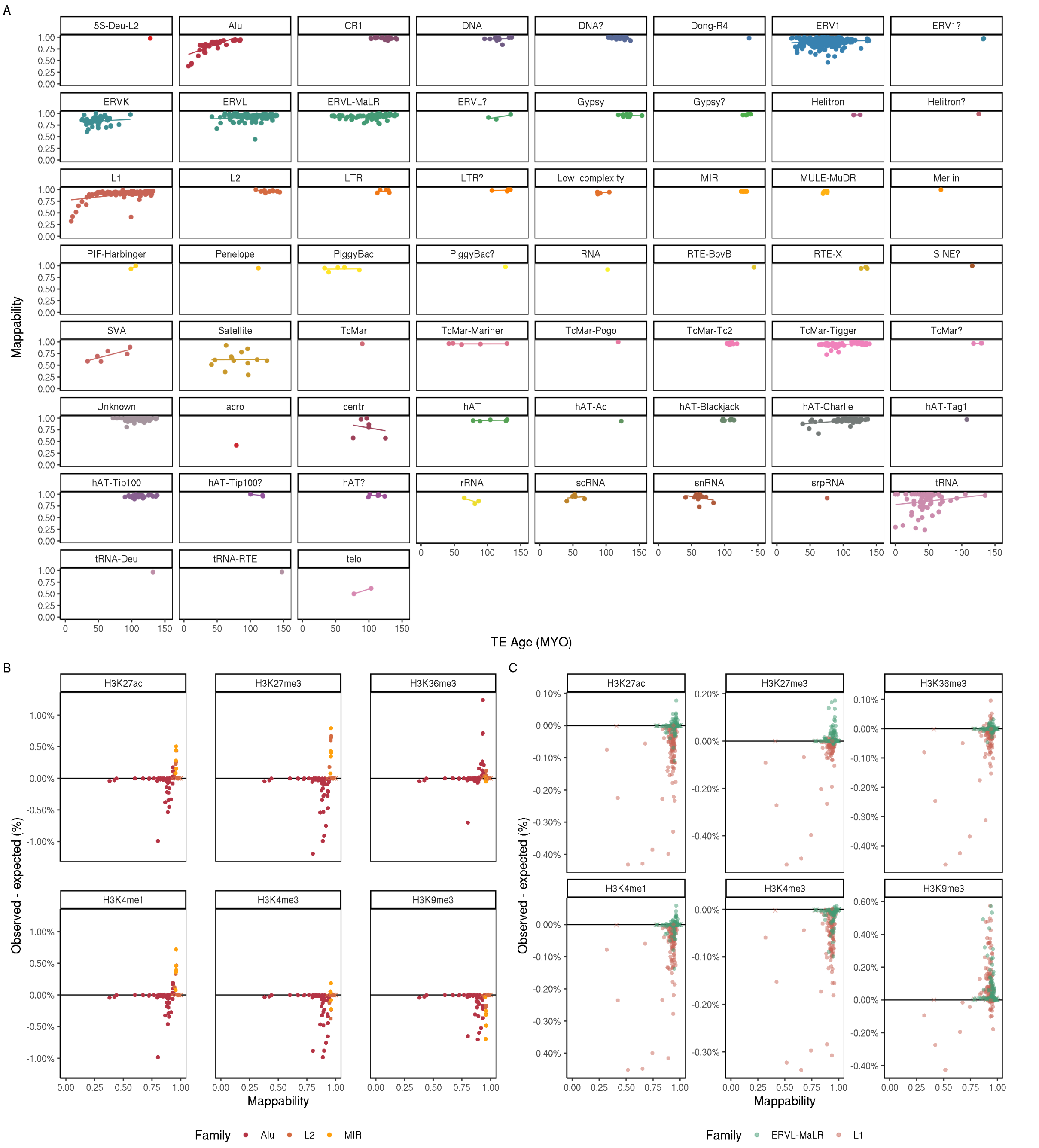


Supplemental Figure 9. TE Mappability and Age. **A)** TE Mappability in function of TE age. **B)** Enrichment in function of mappability for Alu, L2 and MIR **C)** Same as B for ERVL-MaLR and L1 TE families

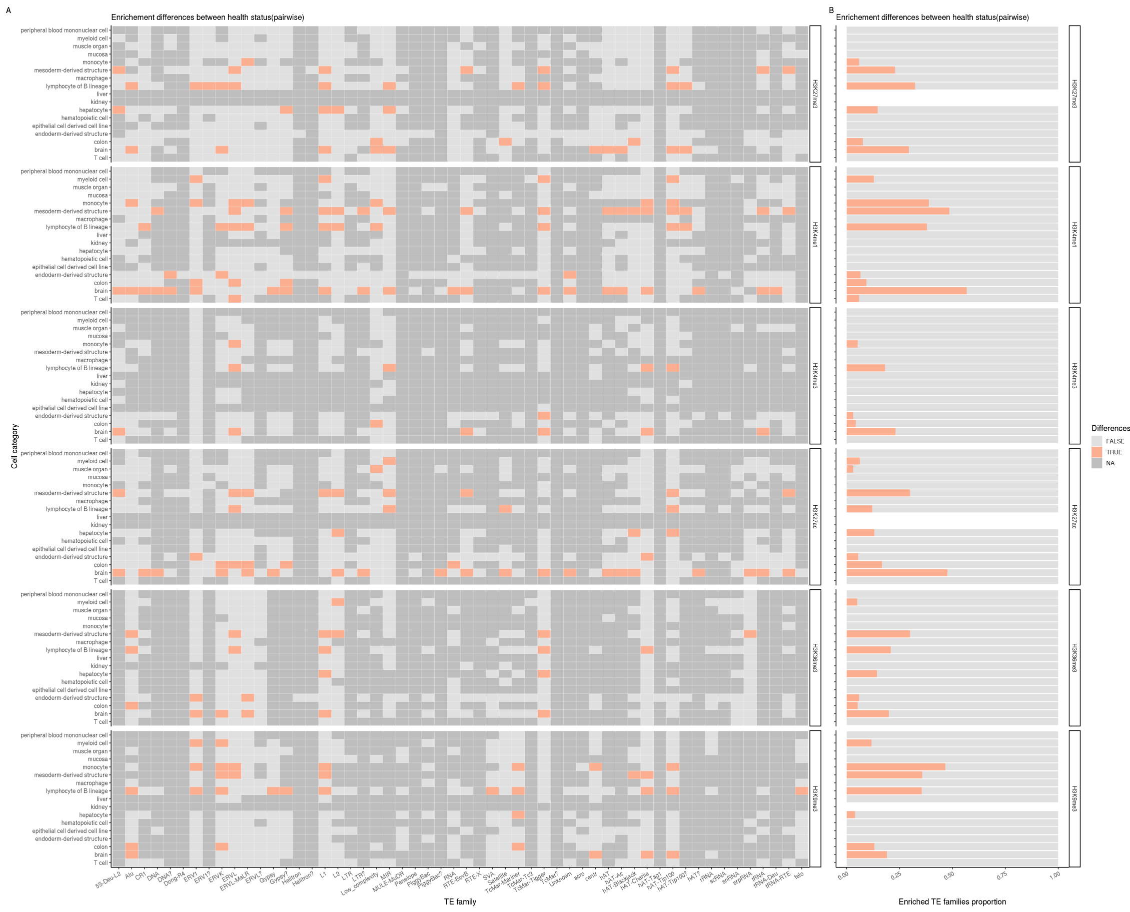


Supplemental Figure 10. Cell type enrichments with significant differences between health statuses. **A)** Comparison between the cell type's health status for the listed TE families. Orange means there was at least 1 significant pairwise difference between the health status, light gray no significant difference, dark gray: Not enough data for the comparison (Only one health status available) **B)** Tally of the proportion TE families with health status differences (orange) or no difference (light gray) for each cell types (Aligned with A).

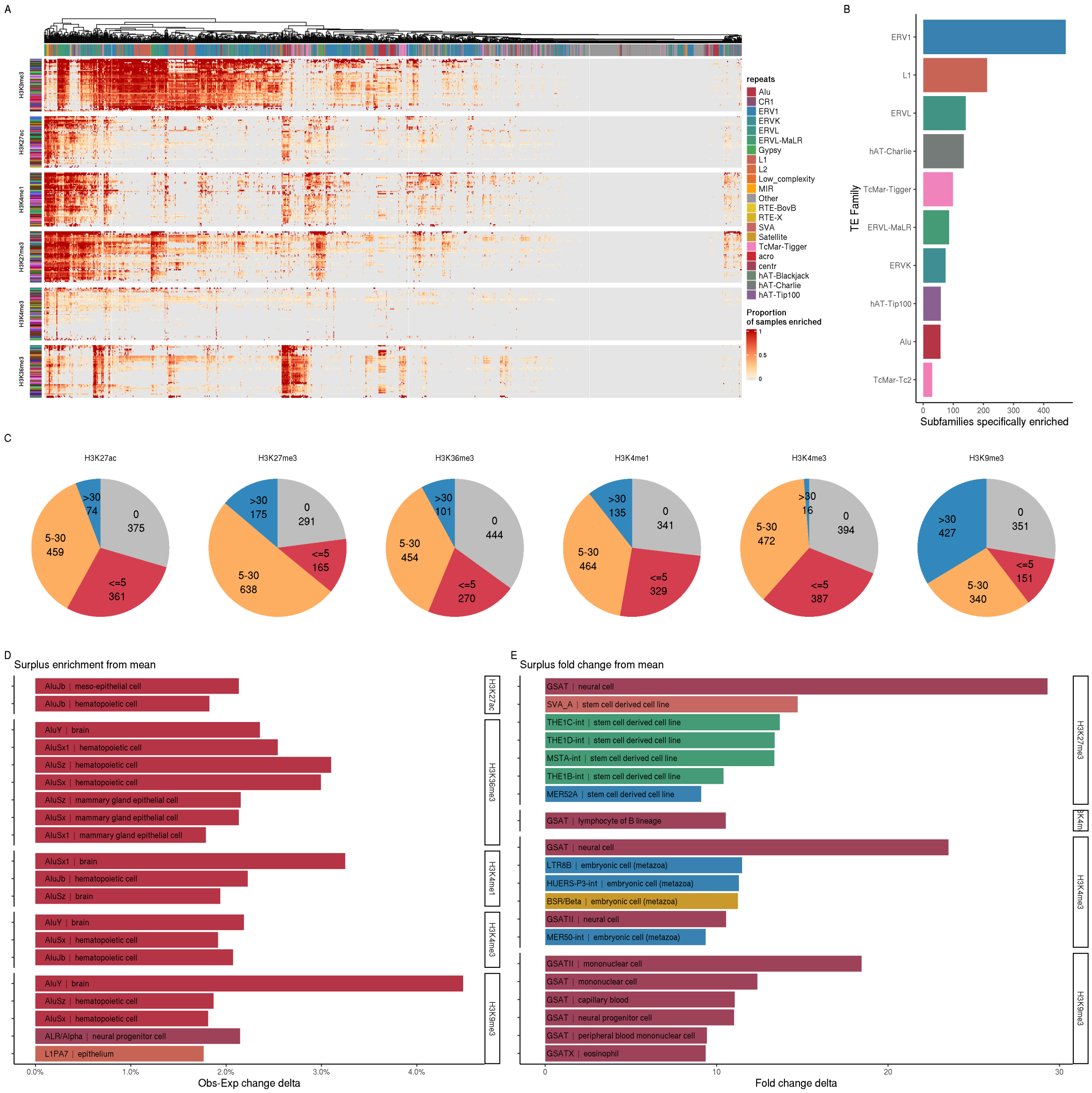


Supplemental Figure 11. Measurements of cell type specificity and surplus **A)** Proportion of samples within each cell type with TE subfamily Enriched (pval<0.001). x axis shows all TE subfamilies, y axis shows the 47 cell types grouped by histone mark. The TE subfamilies are hierarchically clustered. **B)** Top 10 families with most TE subfamilies (with at least 100 instances) enriched in less than 6 cell types (red part of fig 5B pie chart) **C)** Number of TE subfamilies that were enriched in bins of numbers of cell types. Separated by histone mark. **D)** Enrichment surplus of TE subfamily for given cell type. Surplus represents difference between the cell type enrichment and the mean across all cell types. **E)** Same as E for Fold change.

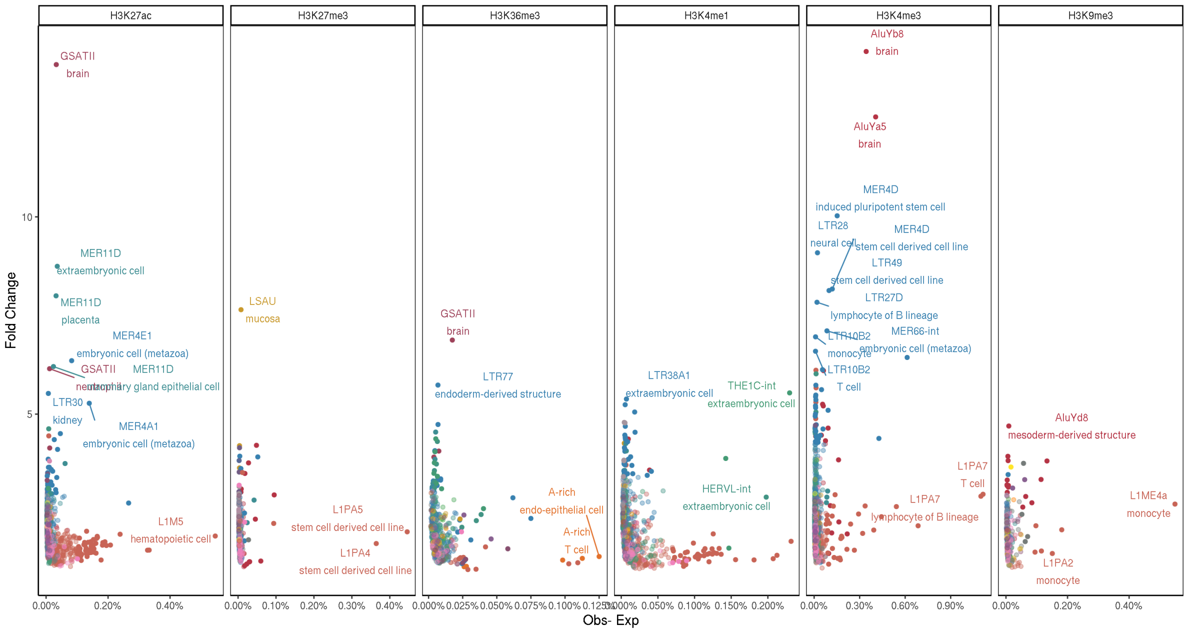


Supplemental Figure 12. Expanded set of Cell type specific TE-histone pairs
Fold change and Observed – Expected TE enrichment of TE subfamilies per cell type. Labeled are random subset of the candidates: most enriched (95th percentile) points in terms of Obs – expected or Fold change.

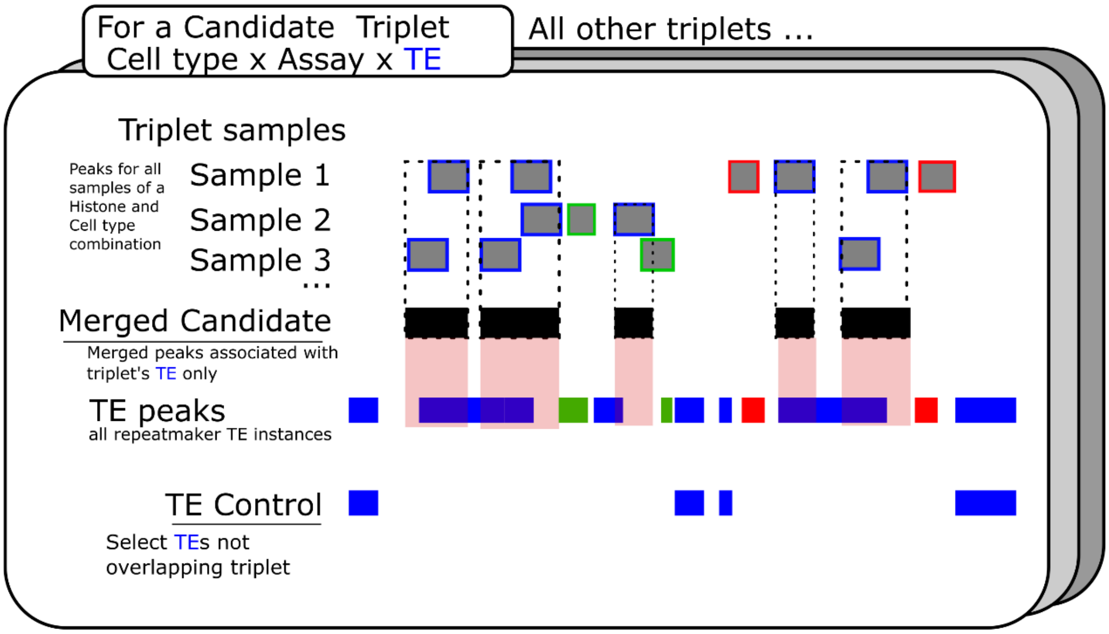


Supplemental Figure 13. Diagram of merged sample and control's generation. (Blue, green and red represent TE families, blue is the TE family of interest for this example) 1. Merge samples from merging all peaks for each Assay-cell type combinations and keeping only the peaks from select TE to compose merged candidates. 2. A TE control keeping only the select TE's instances that were not in the aforementioned triplet. A merged candidate file and associated TE control was generated for all candidates.

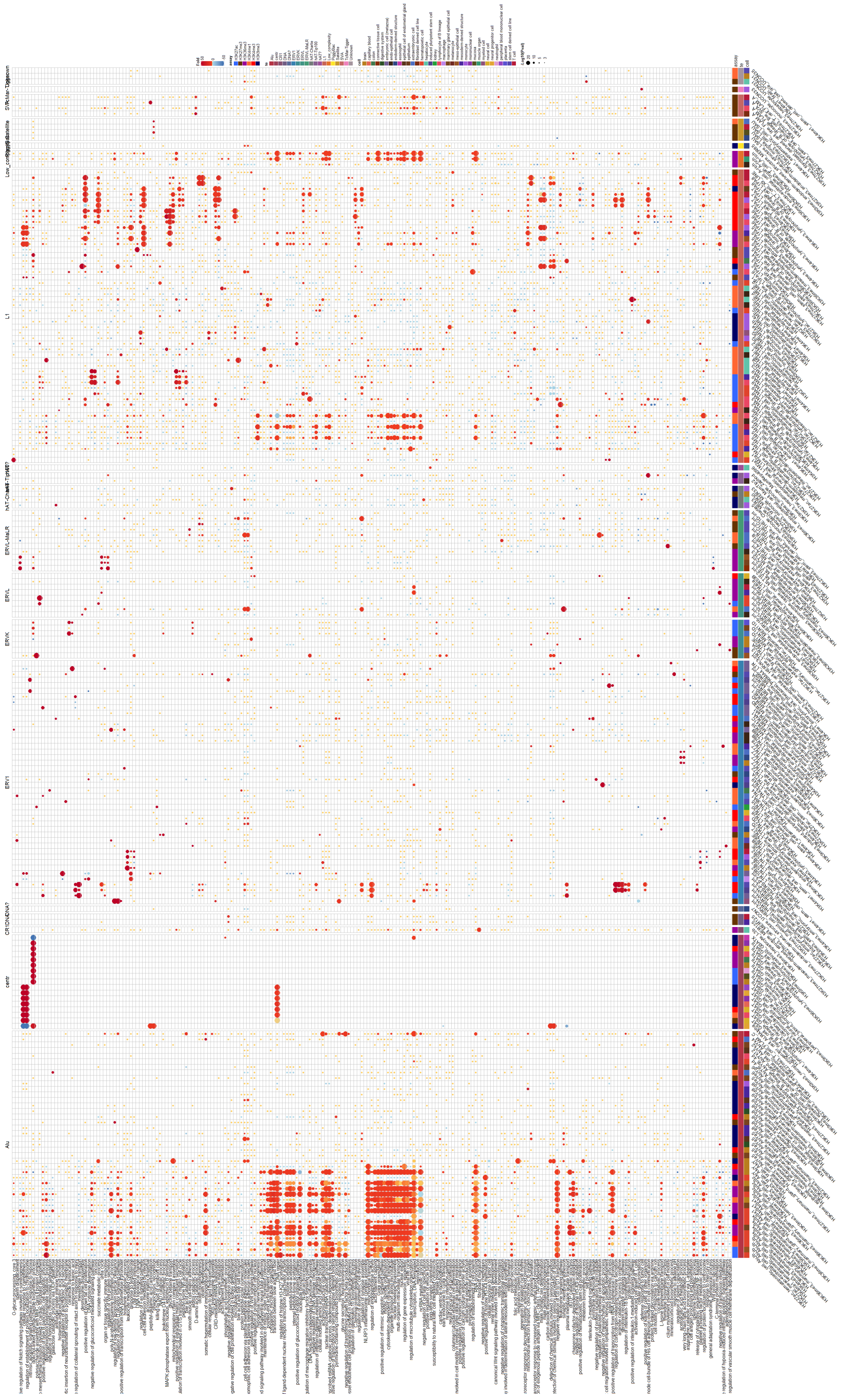
L1
ERVL-MaLR
ERVL
ERVK
ERV1
centr
Alu

Supplemental Figure 14. Fold change enrichment difference from TE control of GO biological processes within for subset of 209 candidate triplet (TE-Assay-Cell Type) samples across histones. (fold change data – fold change of associated TE control) red enriched, yellow between -1 and 1, blue depleted. Circle size represents significance of the enrichment. Terms selected based on data, favored most enriched terms per candidates.

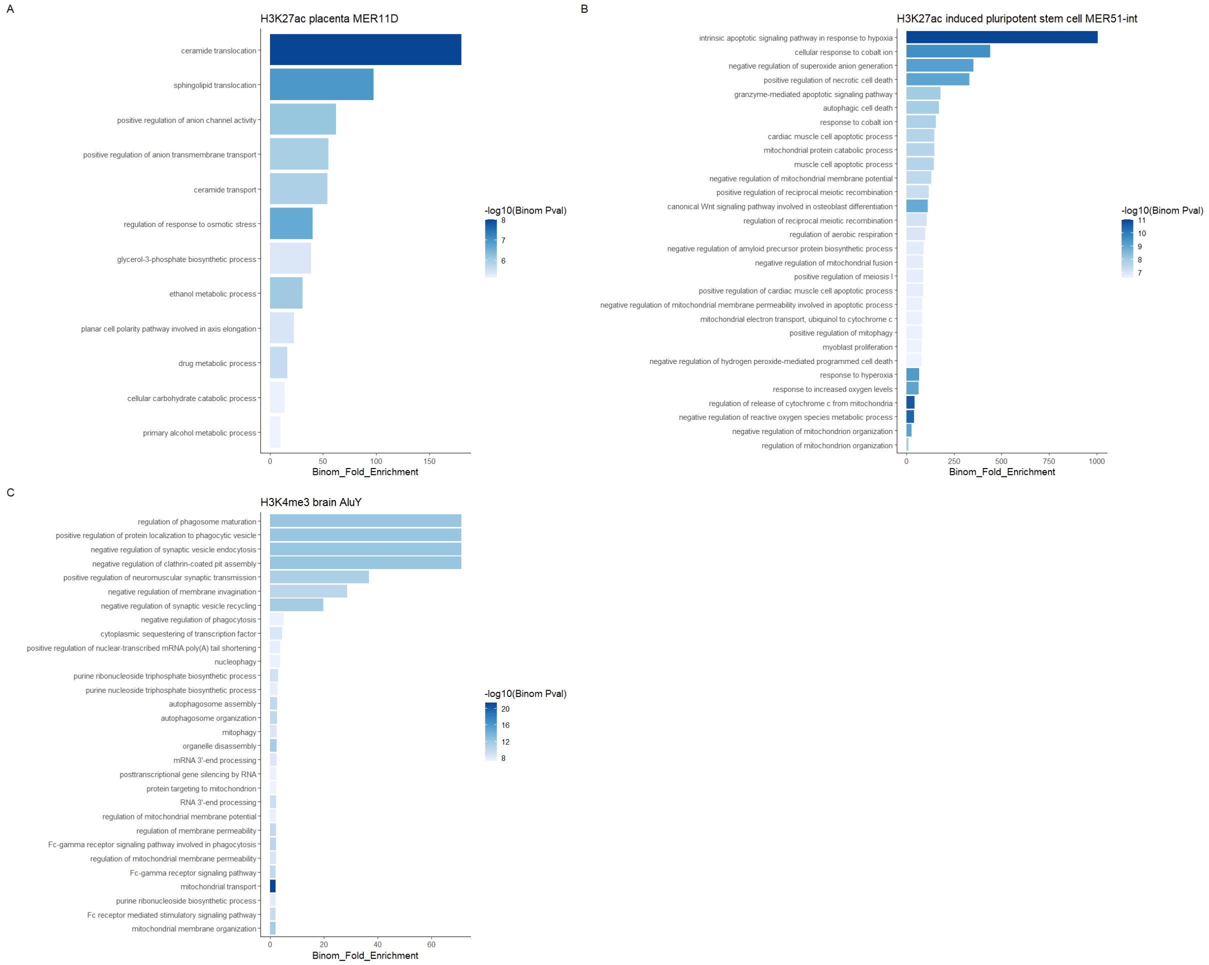


Supplemental Figure 15. Fold change enrichment of GO biological processes within select candidate triplets. **A)** within H3K27ac placenta MER11D triplet. **B)** Within H3K27ac IPS cell MER51-int triplet. **C)** Within H3K4me3 brain AluY. For all plots, showing up to the top 30 processes with fold enrichment >= 2 and –log10(pval) > 5.

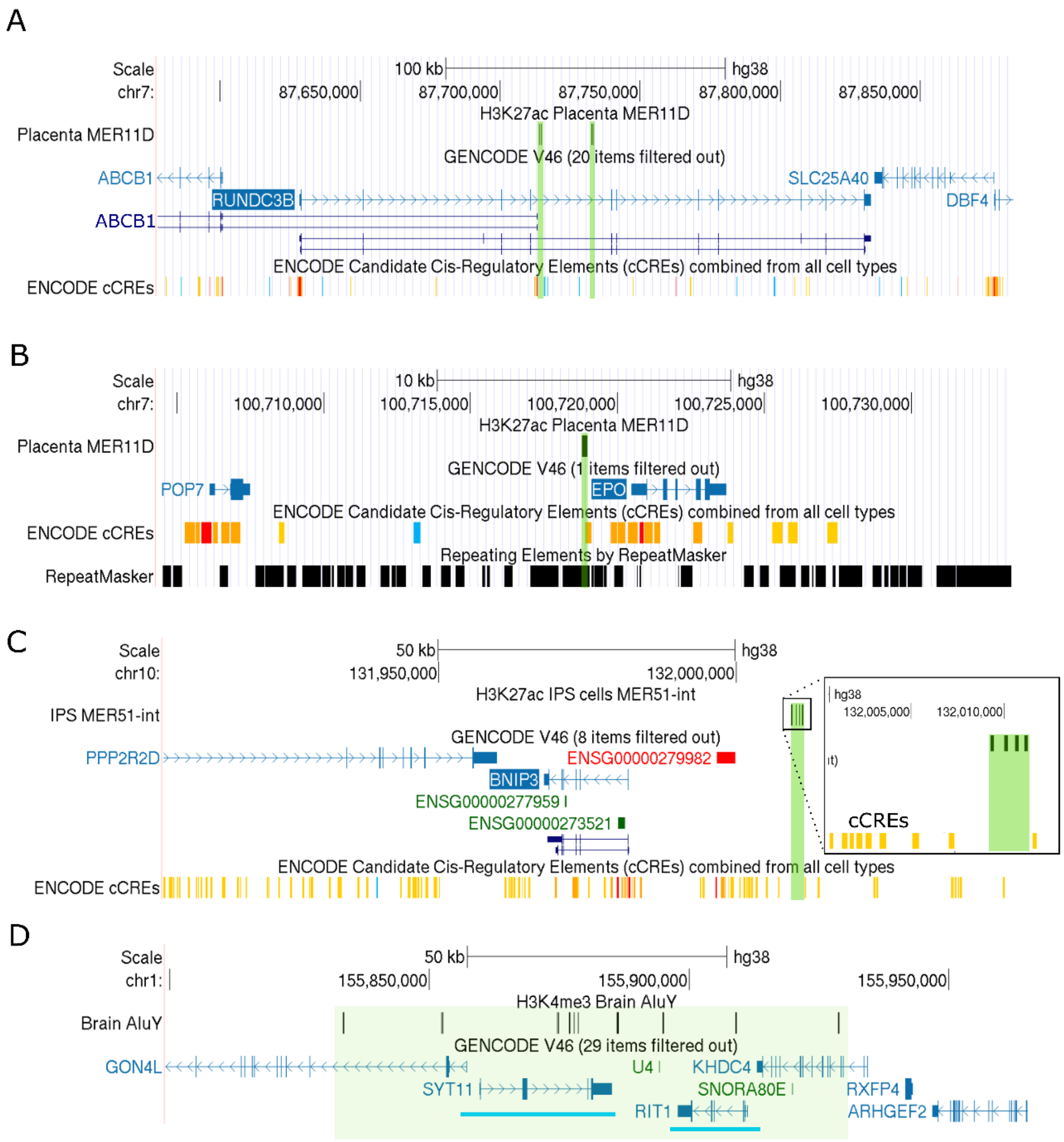


Supplemental Figure 16. Genome tracks of candidate associated peaks. **A)** H3K27ac Placenta peaks overlapping MER11D near ABCB1 gene. **B)** H3K27ac Placenta peak overlapping MER11D near EPO gene. **C)** H3K27ac iPSCs peaks overlapping MER51-int near BNIP3 gene. Close up shows the peaks not overlapping cCREs. **D)** Cluster of H3K4me3 brain peaks overlapping AluY around SYT11 and RIT1 genes.

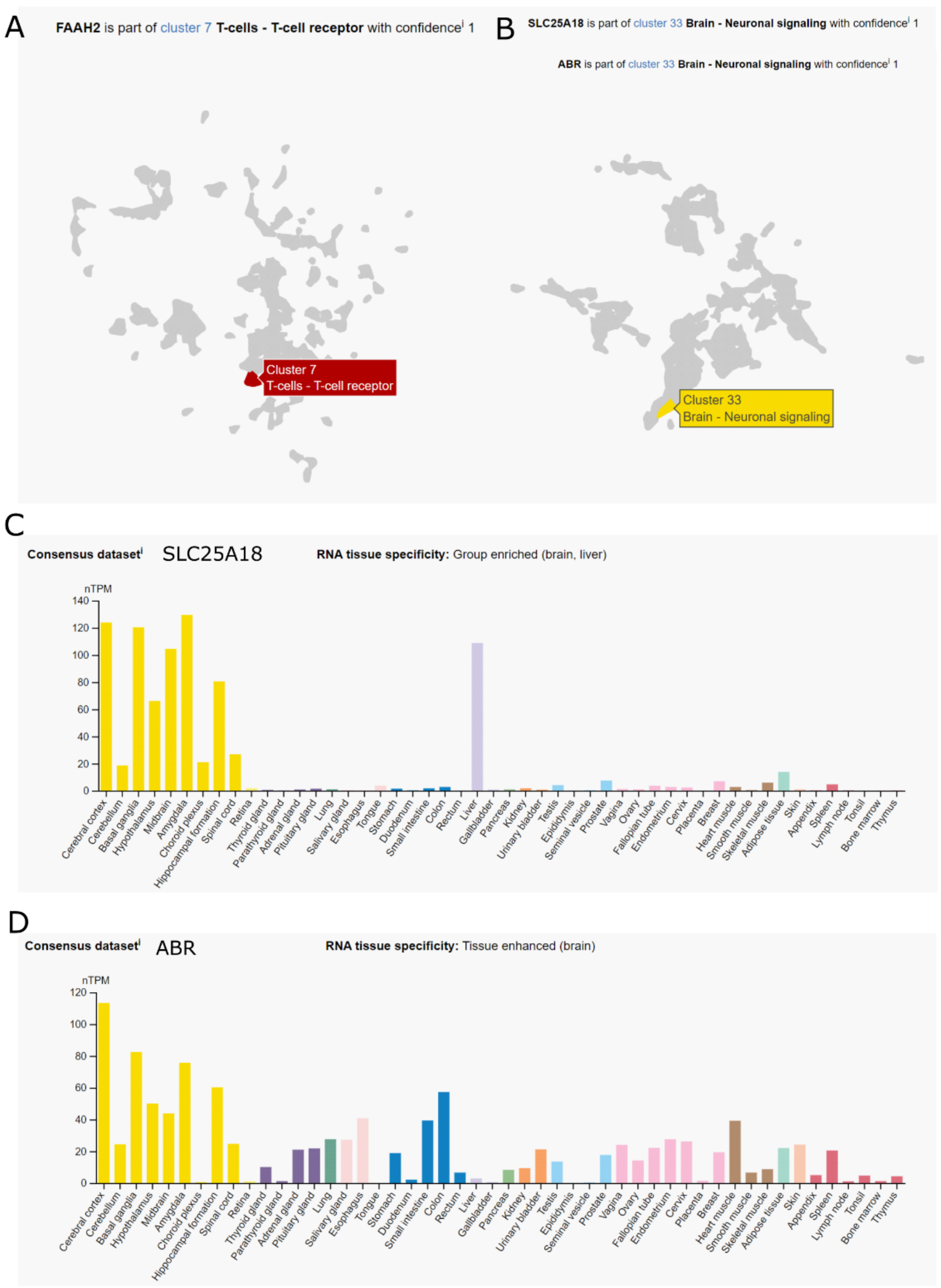


Supplemental Figure 17. RNA tissue expression of select genes. **A)** Single cell expression clustering of FAAH2 gene from protein Atlas(https://www.proteinatlas.org/ENSG00000165591-FAAH2/single+cell+type) **B)** Expression clustering of SCL25A18 and ABR genes, they both grouped in the same cluster. **C)** RNA normalized expression across tissues for SLC15A18 gene(https://www.proteinatlas.org/ENSG00000182902-SLC25A18/tissue) **D)** Same as C for ABR gene(https://www.proteinatlas.org/ENSG00000159842-ABR/tissue). Images and data available from v23.proteinatlas.org

### Supplemental Tables

| |
|---|
| **Supplementary Table 1**.<br>Cell type TE enrichement.<br>Summary statistics (Mean, Median, Max, Min, n) and rank (cell type per assay and TE family) of TE enrichments. Only TE subfamilies that were significantly enriched (Obs-Exp, thus positive) were used and added up for one TE family measurement per sample. The summary statistics are obtained from the summary of all samples grouped by assay and cell type. |
| STable1_summary_te_cell_data_enriched.csv |

| |
|---|
| **Supplementary Table 2**<br>Cell type TE enrichment including non-significant (and depleted) elements<br>Summary statistics (Mean, Median, Max, Min, n) and rank (cell type per assay and TE family) of TE enrichments. All TE subfamilies that were used were used and added up for one TE family measurement per sample. Negative values are possible for cases where Obs-Exp was depleted. The summary statistics are obtained from the summary of all samples grouped by assay and cell type. (same as table 1, but including non significantly enriched TEs) |
| STable2_summary_te_cell_data_all.csv |

| |
|---|
| **Supplementary Table 3**<br>Select candidates table full set<br>Candidate TE name, family and associated assay and cell type with the TE enrichment of that grouping as obs-exp (count_obs_exp_percent) and fold change (count_fold_change). For candidates identified by surplus, the difference from the mean (cell_mean_delta, cell_mean_fold_delta) are also listed. The sample count (n), number of times the observed count was higher than expected (time_over,1000 trial per sample) and resulting pvalue (pval) are listed. And the 4 candidate identifications are shown as true or false (top_obs, top_foldchange, obs_surplus, foldchange_surplus) with the valid method count (method_count) in the last column for each candidate. |
| STable3_TE_candidates_full_set.csv |

| |
|---|
| **Supplementary Table 4**<br>Select candidates table 209 subset<br>Candidate TE name, family and associated assay and cell type with the TE enrichment of that grouping as obs-exp (count_obs_exp_percent) and fold change (count_fold_change). For candidates identified by surplus, the difference from the mean (cell_mean_delta, cell_mean_fold_delta) are also listed. The sample count (n), number of times the observed count was higher than expected (time_over,1000 trial per sample) and resulting pvalue (pval) are listed. And the 4 candidate identifications are shown as true or false (top_obs, top_foldchange, obs_surplus, foldchange_surplus) with the valid method count (method_count) in the last column for each candidate.<br>Subset visualised that was selected as a subset of the top 15 most enriched candidates per histone for top specific obs-exp and fold change using their respective metric (obs-exp and fold change, respectively). And keeping the 40 surplus candidates. |
| STable4_TE_candidates_subset.csv |
